# Cell Types of Origin in the Cell Free Transcriptome in Human Health and Disease

**DOI:** 10.1101/2021.05.05.441859

**Authors:** Sevahn K. Vorperian, Mira N. Moufarrej, Tabula Sapiens Consortium, Stephen R. Quake

## Abstract

Cell-free RNA (cfRNA) can be used to noninvasively measure dynamic and longitudinal physiological changes throughout the body. While there is considerable effort in the liquid biopsy field to determine disease tissue-of-origin, pathophysiology occurs at the cellular level. Here, we describe two approaches to identify cell type contributions to cfRNA. First we used *Tabula Sapiens*, a transcriptomic cell atlas of the human body to computationally deconvolve the cell-free transcriptome into a sum of cell type specific transcriptomes, thus revealing the spectrum of cell types readily detectable in the blood. Second, we used individual tissue transcriptomic cell atlases in combination with the Human Protein Atlas RNA consensus dataset to create cell type signature scores which can be used to infer the implicated cell types from cfRNA for a variety of diseases. Taken together, these results demonstrate that cfRNA reflects cellular contributions in health and disease from diverse cell types, potentially enabling determination of pathophysiological changes of many cell types from a single blood test.

## Main Text

Cell-free RNA (cfRNA) represents a mixture of transcripts reflecting the health status of multiple tissues, thereby affording broad clinical utility. Existing applications span oncology and bone marrow transplantation^12^, obstetrics^3,45^, neurodegeneration^6^, and liver disease^7^. However, several aspects about the physiologic origins of cfRNA including the contributing cell types-of-origin remain unknown, and current assays focus on tissue level contributions at-best^2,4,6–8^. Incorporating knowledge from cellular pathophysiology, which often forms the basis of disease^9^, into a liquid biopsy would more closely match the resolution afforded by invasive procedures.

We first characterized the landscape of cell type specific signal from healthy donor plasma (Fig. 1A) using published exome-enriched cf-transcriptome data^6^ (Fig. 1A). After removing low-quality samples (Fig. S1, Methods), we intersected the set of genes detected in healthy individuals (n = 75) with a database of cell-type specific markers defined in context of the whole body^10^. Marker genes for blood, brain, and liver cell types were readily detected, as previously observed at tissue level^1,4,6,7 2^as well as the kidney, gastrointestinal track, and pancreas (Fig. 1B).

**Fig. 1:**
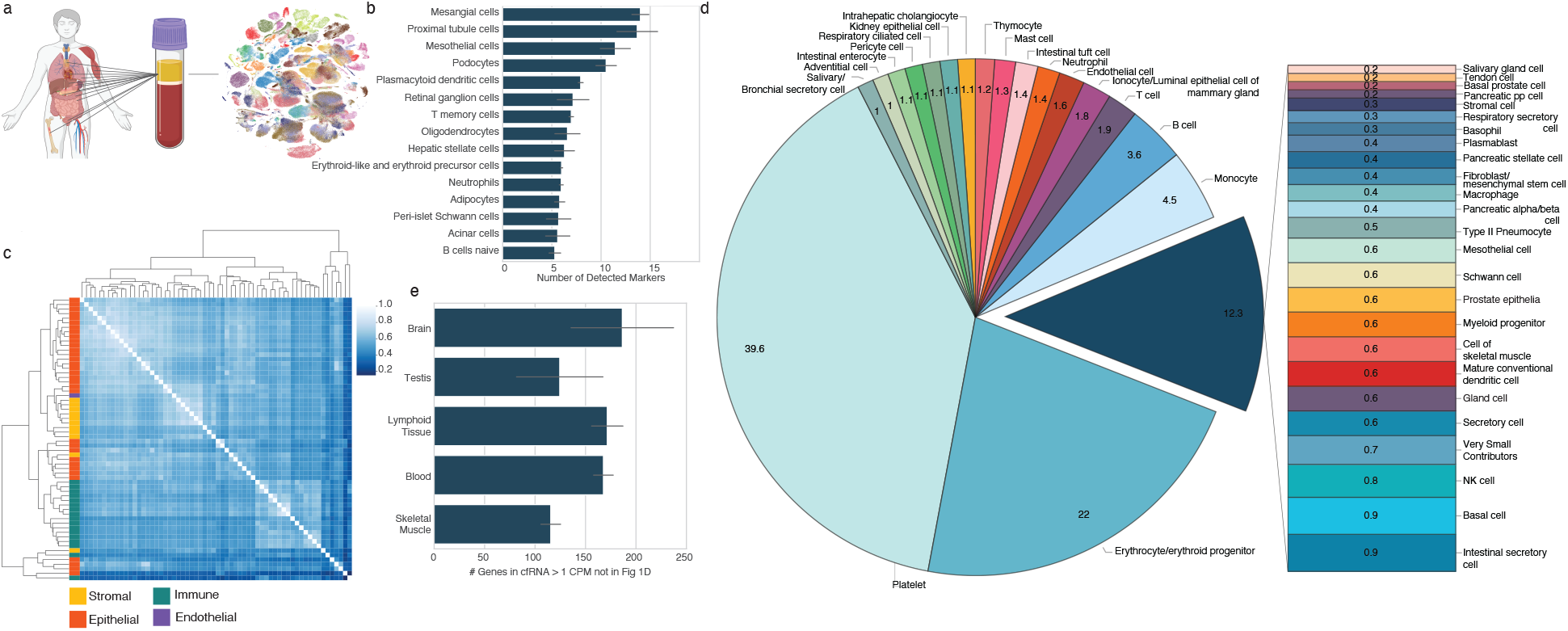
Cell type decomposition of the plasma cell free transcriptome using Tabula Sapiens reveals comprehensive cell type-of-origin landscape. a. Schematic depicting integration of tissue-of-origin and single cell transcriptomics to identify cell types-of-origin in plasma cfRNA. b. Number of cell type specific markers defined in context of the human body that are identified in cfRNA. Bars and error bars respectively denote the mean and s.d. of number of cell type specific markers across patients (n = 75). CPM-TMM counts for a given gene across technical replicates were averaged prior to intersection. c. Cluster heatmap of Spearman correlations of cell type basis matrix column space derived from Tabula Sapiens. d. Mean fractional contributions of cell type specific RNA in plasma cf-transcriptome (n = 18). e. Top tissues in cfRNA not captured by basis matrix (e.g. the set difference of all genes detected in a given cfRNA sample and the row space of the basis matrix intersection with HPA tissue specific genes). Bars and error bars respectively denote the mean and s.d. of number of HPA tissue specific genes with NX counts > 10 and cell free CPM-TMM normalized expression ≥ 1 (n = 18).

We then sought to deconvolve the fractions of cell-type specific RNA using support vector regression, a deconvolution method previously applied to decompose bulk tissue transcriptomes into fractional cell type contributions^11,12^. We used *Tabula Sapiens* v1.0 (TSP)^13^, a multiple donor whole body cell atlas spanning 24 tissues and organs, to define a basis matrix whose gene set accurately and simultaneously resolved the distinct cell types in TSP. The basis matrix was defined using the gene space that maximized linear independence of the cell types; and does not include the whole transcriptome but rather the minimum discriminatory gene set to distinguish between the cell types in Tabula Sapiens. To reduce multicollinearity, transcriptionally similar cell types were grouped (fig. S2). We observed that the basis matrix defined by this gene set appropriately described cell types as most similar to others from the same organ compartment and correspond to the highest off-diagonal similarity (Fig. 1C). We also confirmed that the basis matrix accurately deconvolved cell type specific RNA fractional contributions from several bulk tissue samples (fig. S3, S4, Supplementary Note 1).

We used this matrix to deconvolve the cell types-of-origin in the plasma cf-transcriptome (Fig. 1D; fig. S5, S6). Platelets, erythrocyte/erythroid progenitors, and leukocytes comprised the majority of observed signal, whose respective proportions were generally consistent with recent estimates from serum cfRNA^1^ and plasma cfDNA^14^. Within this set of cell types, we suspect that the observation of platelets as a majority cell type, rather than megakaryocytes^1^, likely reflects annotation differences in reference data. We observed distinct transcriptional contributions from solid tissue-specific cell types from the intestine, liver, lungs, pancreas, heart, and kidney (Fig. 1D, fig. S5). Altogether, the observation of contributions from many non-hematopoietic cell types underscores the ability to simultaneously noninvasively resolve contributions to cfRNA from disparate cell types across the body.

Some cell types likely present in the plasma cf-transcriptome were missing in this decomposition because the source tissues were not represented in TSP. Though ideally reference gene profiles for all cell types would be simultaneously considered in this decomposition, a complete reference dataset spanning the entire cell type space of the human body does not yet exist. To identify cell type contributions possibly absent from this analysis, we intersected the genes measured in cfRNA absent from the basis matrix with tissue specific genes from the human protein atlas RNA consensus dataset (HPA)^15^. This identified both the brain and testes as tissues whose cell types were not found during systems-level deconvolution and additional genes specific to the liver, skeletal muscle, and lymphoid tissues that were not used by the basis matrix (Fig. 1E, Methods).

As an example of how to analyze cell type contributions from tissues which were not present in TSP, we utilized an independent brain single cell atlas along with HPA to define cell type gene profiles and examined their expression in cfRNA (Fig 2A, fig. S7–9). There was a strong signature score from excitatory neurons and a reduced signature score from inhibitory neurons. We observed strong signals from astrocytes, oligodendrocytes, and oligodendrocyte precursor cells. These glial cells facilitate brain homeostasis, form myelin, and provide neuronal structure and support^9^, consistent with evidence of RNA transport across and the permeability of the blood brain barrier^16,17^ and that some brain regions are in direct contact with the blood ^18^. Similarly, we utilized published cell atlases for the placenta^19,20^, kidney^21^, and liver^22^ to define cell type specific gene profiles (fig. S8, S10) from which to perform signature scoring. These observations augment the resolution of previously observed tissue-specific genes reported to date in cfRNA^4,6,7 21^and formed a baseline from which to measure aberrations in disease.

**Fig. 2:**
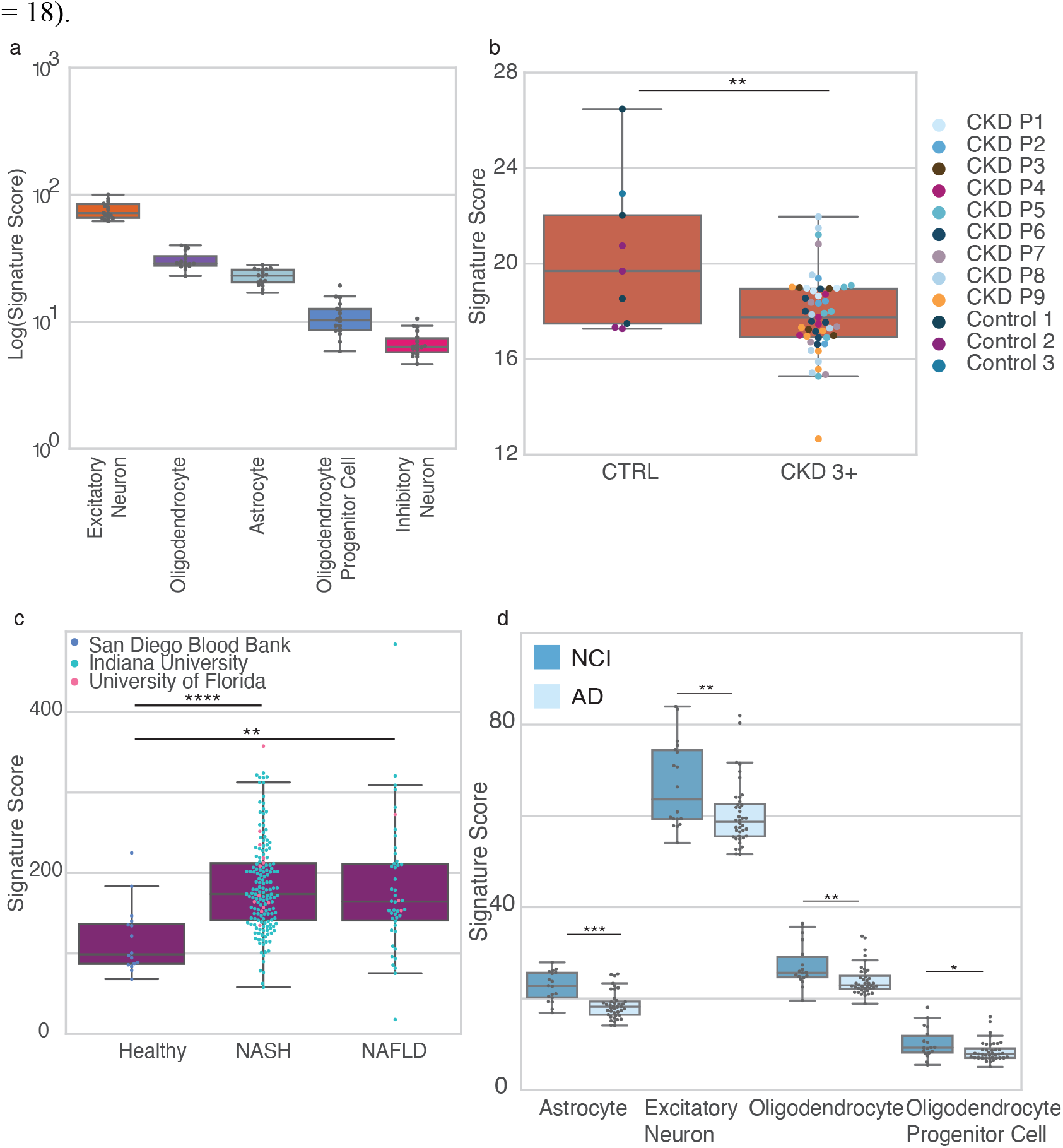
Cellular pathophysiology is noninvasively resolvable in cfRNA. For a given boxplot, any cell type signature score is the sum of log-transformed CPM-TMM normalized counts. Box plot: horizontal line, median; lower hinge, 25^th^ percentile; upper hinge, 75^th^ percentile; whiskers,1.5 interquartile range; points outside whiskers indicate outliers. All P values were determined by a Mann Whitney U test; sidedness specified in subplot caption. \**P* < 0.05, \*\**P* < 10^−2^, \*\*\**P* < 10^−4^, \*\*\*\**P* < 10^−5^. a. Neuronal and glial cell type signature scores in healthy cfRNA plasma (n = 18) on a logarithmic scale. b. Comparison of dimensionless signature scores CKD stages 3+ (n = 51 samples; 9 patients) and healthy controls (n = 9 samples; 3 patients) using data from Ibarra et al. Dot color denotes each patient (*P* = 9.66*10^−3^, U = 116, one-sided) c. Hepatocyte signature scure between healthy (n = 16) and both NAFLD (n = 46) (*P* =3.15 x 10^−4^, U = 155, one-sided) and NASH (n = 163) (*P* = 4.68 x 10^−6^, U = 427, one-sided); NASH vs. NAFLD (*P* = 0.464, U = 3483, two-sided). Color reflects sample collection center. d. Neuronal and glial signature scores in AD (n = 40) and NCI (n = 18) cohorts. Excitatory neuron (*P* = 4.94 x 10^−3^, U = 206, one-sided), oligodendrocyte (*P* = 2.28 x 10^−3^, U = 178, two-sided), oligodendrocyte progenitor (*P* = 2.27 x 10^−2^, U = 224, two-sided), and astrocyte (*P* = 6.1 x 10^−5^, U = 121, two-sided).

Cell type specific changes drive disease etiology^9^, and we asked whether cfRNA reflected cellular pathophysiology. We considered trophoblasts in preeclampsia^23,24^, proximal tubules in chronic kidney disease (CKD)^25,26^, hepatocytes in non-alcoholic steatohepatitis (NASH)/non-alcoholic fatty liver disease (NAFLD)^27^, and multiple brain cell types in Alzheimer’s Disease (AD)^28,29^. As an example of why whole body cell type characterization is important, we observed that a prior attempt to infer trophoblast cell types from cfRNA in preeclampsia^24^ used genes that are not specific or readily measurable within their asserted cell type (fig. S11, Supplementary Note 2). However, we found several other cases where cellular pathophysiology can be measured in cfRNA.

The proximal tubule is a highly metabolic, predominant kidney cell type and is a major source for injury and disease progression in CKD^25,26^. Tubular atrophy is a hallmark of CKD nearly independent of disease etiology^30^ and is superior to clinical gold standard as a predictor of CKD progression^31^. Using data from Ibarra et al, we discovered a striking decrease in the proximal tubule cell signature score of CKD patients (ages 67-91, CKD stage 3-5 or peritoneal dialysis) compared to healthy controls (Fig. 2B, fjg. S12). These results demonstrate noninvasive resolution of proximal tubule deterioration observed in CKD histology^31^ and are consistent with findings from invasive biopsy.

Hepatocyte steatosis is a histologic hallmark of NASH and NAFLD phenotypes, whereby the accumulation of cellular stressors results in hepatocyte death^27^. We found that several genes differentially expressed in NAFLD serum cfRNA^7^ were specific to the hepatocyte cell type profile derived above. (*P* < 10^−10^, hypergeometric test). Notable hepatocyte specific DEG include cytochrome P450 enzymes (including *CYP1A2, CYP2E1*, *CYP3A4*), lipid secretion (*MTTP)*, and hepatokine-coding genes (*AHSG, LECT2)*^32^. We further observed striking differences in the hepatocyte signature score between healthy and both NAFLD and NASH (Fig. 2C, fig. S13), and no difference between the NASH and NAFLD cohorts.

AD pathogenesis results in neuronal death and synaptic loss^29^. We used brain single cell data^28^ to define brain cell type gene profiles in both the AD and non-cognitively impaired (NCI) brain. Many of the differentially expressed genes found in cfRNA analysis of AD plasma are brain cell type specific (*P* < 10^−5^, hypergeometric test). Astrocyte specific genes included filament protein (*GFAP*^33^) and ion channels (*GRIN2C* ^28^). Excitatory neuron specific genes include solute carrier proteins (*SLC17A7*^28^ & *SLC8A2*^34^), cadherin proteins (*CDH8*^35^ & *CDH22*^36^), and a glutamate receptor (*GRM1*^29,37^). Oligodendrocyte-specific genes encode proteins for myelin sheath stabilization (*MOBP*^29^) and a synaptic/axonal membrane protein (*CNTN2*^29^). Oligodendrocyte precursor cell-specific genes included transcription factors (*OLIG2*^38^ & *MYT1*^39^), neural growth and differentiation factor (*CSPG5*^40^), and a protein putatively involved in brain extracellular matrix formation (*BCAN*^41^).

We then inferred neuronal death between AD and NCI and also observed differences in oligodendrocyte, oligodendrocyte progenitor, and astrocyte signature scores (Fig. 2D, fig. S14). The oligodendrocyte and oligodendrocyte progenitor cells signature score directionality agrees with reports of their death and inhibited proliferation in AD respectively^42^. The observed astrocyte signature score directionality is consistent with the cell type specificity of a subset of reported downregulated DEG^6^ and reflects that astrocyte-specific changes, which are known to in AD pathology^42^, are noninvasively measurable.

Taken together, this work demonstrates consistent noninvasive detection of cell-type specific changes in human health and disease using cfRNA. Our findings uphold and further augment the scope of prior work identifying immune cell types^1^ and hematopoietic tissues^1,4^ as primary contributors to the cf-transcriptome cell type landscape. Our approach is complementary to prior work using cell free nucleosomes^14^ which depends on a more limited set of reference ChIP-seq data which is largely at the tissue level^43^. Readily measurable cell types include those specific to the brain, lung, intestine, liver, and kidney, whose pathophysiology affords broad prognostic and clinical importance. Consistent detection of cell types responsible for drug metabolism (e.g. liver and renal cell types) as well as cell types that are drug targets, such as neurons or oligodendrocytes for Alzheimer’s-protective drugs, could provide strong clinical trial end-point data when evaluating drug toxicity and efficacy. We anticipate that the ability to noninvasively resolve cell type signatures in plasma cfRNA will both enhance existing clinical knowledge and enable increased resolution in monitoring disease progression and drug response.

## Acknowledgements

We thank M. Chen for single cell analysis input, feedback and helpful discussions. We thank E. Sattely and G.E. Marti for helpful discussions. We thank G. Loeb for kidney discussions. The human body in figure 1A was created using Biorender.com.

## Author contributions

S.K.V. and S.R.Q. conceptualized the study. S.K.V. and S.R.Q. designed the study in collaboration with M.N.M. S.K.V. performed all analyses; M.N.M. wrote the bioinformatic preprocessing pipeline to map reads to the human genome and cell free sample QC. S.K.V, M.N.M, S.R.Q wrote the manuscript. All authors revised the manuscript and approved it for publication.

## Data availability

All datasets used for this work were publicly available, downloaded with permission, or directly requested from authors. Tissue gene lists and NX counts were downloaded from HPA (www.proteinatlas.org, v19). GTEx raw expression was downloaded from the GTEx portal (https://www.gtexportal.org/home/datasets, GTEx analysis V8). Tabula Sapiens single cell data were received from the CZ-Biohub (https://tabula-sapiens-portal.ds.czbiohub.org, version 1.0). The brain single cell data were downloaded with permission from Synapse (https://www.synapse.org/#!Synapse:syn18485175) and associated ROSMAP metadata were downloaded with permission from Synapse (https://www.synapse.org/#!Synapse:syn3157322). The liver Seurat object was requested from Aizarani et al. For the placenta atlases, a Seurat object was requested from Suryawanshi et al and AnnData requested from Vento-Tormo et al. Kidney AnnData was downloaded (https://www.kidneycellatlas.org, Mature Full dataset).

## Code availability

Code for the work in this manuscript will be available on Github at www.github.com/sevahn/deconvolution.

## Materials & Methods

### Data Processing

#### Data acquisition

Cell free RNA: For samples from Ibarra et al, raw sequencing data was obtained from the SRA (PRJNA517339). For samples from Munchel et al, processed counts tables were directly downloaded.

For all individual tissue single cell atlases, Seurat objects or AnnData objects were downloaded or directly received from authors. Data from Mathys et al. were downloaded with permission from Synapse.

HPA v19 transcriptomic data, GTEx v8 raw counts, and Tabula Sapiens v1.0 were downloaded directly.

#### Bioinformatic processing

For each sample for which raw sequencing data were downloaded, we trimmed reads using trimmomatic (v 0.36) and then mapped them to the human reference genome (hg38) with STAR (v 2.7.3a). Duplicate reads were then marked and removed by GATK’s (v 4.1.1) MarkDuplicates tool. Finally, mapped reads were quantified using htseq-count (v 0.11.1), and read statistics were estimated using FastQC (v 0.11.8).

The bioinformatic pipeline was managed using snakemake (v 5.8.1). Read and tool performance statistics were aggregated using MultiQC (v 1.7).

#### Sample quality filtering

For every sample for which raw sequencing data was available, we estimated three quality parameters as previously described^44,45^: RNA degradation, ribosomal read fraction, and DNA contamination. RNA degredation was estimated by calculating a 3’ bias ratio. Specifically, we first counted the number of reads per exon and then annotated each exon with its corresponding gene ID and exon number using htseq-count. Using these annotations, we measured the frequency of genes for which all reads mapped exclusively to the 3’ most exon as compared to the total number of genes detected. We approximate RNA degradation for a given sample as the fraction of genes where all reads mapped to the 3’ most exon. To estimate ribosomal read fraction, we compared the number of reads that mapped to the ribosome (Region GL00220.1:105424-118780, hg38) relative to the total number of reads (Samtools view). To estimate DNA contamination, we used an intron to exon ratio and quantified the number of reads that mapped to intronic as compared to exonic regions of the genome.

We applied the following thresholds as previously reported^44^:

- Ribosomal: > 0.2
- 3’ Degradation: > 0.4
- Intron/Exon: > 3

We considered any given sample as low quality if its value for any metric was greater than any of these thresholds and excluded the sample from subsequent analysis.

#### Data Normalization

All gene counts were adjusted to counts per million reads and per milliliter of plasma used. For a given sample (*i* denotes gene index and *j* denotes sample index):

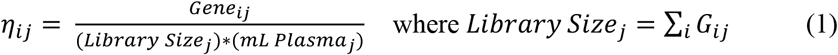

For subjects who had samples with multiple technical replicates, these plasma volume CPM counts were averaged prior to nu-SVR deconvolution.

For all analyses except nu-SVR (e.g. all work except Fig. 1D and 1E), we next applied trimmed mean of M values (TMM) normalization as previously described^46^:

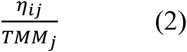

CPM TMM normalized gene counts across technical replicates for a given biological replicate were averaged for the count tables used in all analyses performed.

Sequencing batches and plasma volumes were obtained from the authors in Toden et al and Chalasani et al for per-sample normalization. For samples from Ibarra et al., plasma volume was assumed to be constant at 1 mL as we were unable to attain this information from the authors; sequencing batches were confirmed with authors (personal communication). All samples from Munchel et al were used to compute TMM scaling factors and 4.5 mL plasma^5^ was used to normalize all samples within a given dataset (both PEARL-PEC and iPEC).

### Cell Type Marker Identification using PanglaoDB

The PanglaoDB cell type marker database was downloaded on March 27, 2020. Markers were filtered for human (“Hs”) only and for PanglaoDB’s defined specificity (how often marker was not expressed in a given cell type) and sensitivity (how frequently marker is expressed in cells of this type). Gene synonyms from Panglao were determined using MyGene version 3.1.0 to ensure full gene space.

We then intersected this gene space with a cohort of healthy cfRNA samples (n = 74, non-cognitively impaired (NCI) individuals from Toden et al). A given cell type marker was counted in a given healthy cfRNA sample if its gene expression was greater than zero in log + 1 transformed CPM-TMM gene count space.

Cell types with markers filtered by sensitivity = 0.9 and specificity = 0.2 and samples with ≥ 5 cell type markers are shown in Fig 1B.

### Basis Matrix Formation

Scanpy^47^ (version 1.6.0) was used. Only cells from droplet sequencing (“10X”) were used in analysis given that a more comprehensive set of unique cell types across the tissues in *Tabula Sapiens* were available^13^. Disassociation genes as reported^13^ were eliminated from the gene space prior to subsequent analysis.

Given the non-specificity of the following annotations (e.g. other cell type annotations at finer resolution existed), cells with these annotations were excluded from subsequent analysis:

- ‘epithelial cell’
- ‘ocular surface cell’
- ’radial glial cell’
- ’lacrimal gland functional unit cell’
- ’connective tissue cell’
- ’corneal keratocyte’
- ’ciliary body’
- ’bronchial smooth muscle cell’
- ’fast muscle cell’
- ’muscle cell’
- ’myometrial cell’
- ’skeletal muscle satellite stem cell’
- ’slow muscle cell’
- ’tongue muscle cell’
- ’vascular associated smooth muscle cell’
- ’alveolar fibroblast’
- ’fibroblast of breast’
- ’fibroblast of cardiac tissue’
- ’myofibroblast cell’

All additional cells belonging to the ‘Eye’ tissue were excluded from subsequent analysis given discrepancies in compartment and cell type annotations and the unlikelihood of detecting eye-specific cell types. The resulting cell type space still possessed several transcriptionally similar cell types (e.g. various intestinal enterocytes, t cells, or dendritic cells), that left unaddressed, would reduce the linear independence of the basis matrix column space and hence would impact nu-SVR deconvolution.

Cells were therefore assigned broader annotations on a per-compartment basis as follows<u>: Epithelial, Stromal, Endothelial:</u> using counts from the ‘decontXcounts’ layer of the adata object, cells were CPM normalized (sc.pp.normalize_total(target_sum = 1e6) and log-transformed (sc.pp.log1p). Hierarchical clustering with complete linkage (sc.tl.dendrogram) was performed per compartment on the feature space comprising the first 50 principal components (sc.pp.pca). Epithelial and stromal compartment dendrograms were then cut (scipy.cluster.hierarchy.cut_tree) at 20% and 10% of the height of the highest node respectively, such that cell types with high transcriptional similarity were grouped together but overall granularity of the cell type labels was preserved. This work is available in the script “cuttree.ipynb” on Github.

The endothelial compartment dendrogram revealed high transcriptional similarity across all cell types (maximum node height = 0.851) compared to epithelial (maximum node height = 3.78) and stromal (maximum node height = 2.34) compartments (fig. S2). To this end, only the ‘endothelial cell’ annotation was utilized for the ‘endothelial’ compartment.

<u>Immune:</u> given the high transcriptional similarity and the varying degree of annotation granularity across tissues and cell types, cell types were grouped on the basis of annotation. The following immune annotations were kept:

- ’b cell’
- ’basophil’
- ’erythrocyte’
- ’erythroid progenitor’
- ’hematopoietic stem cell’
- ’innate lymphoid cell’
- ’macrophage’
- ’mast cell’
- ’mature conventional dendritic cell’
- ’microglial cell’
- ’monocyte’
- ’myeloid progenitor’
- ’neutrophil’
- ’nk cell’
- ’plasma cell’
- ’plasmablast’
- ’platelet’
- ’t cell’
- ’thymocyte’

All other immune compartment cell type annotations were excluded for being too broad when more detailed annotations existed (i.e. ‘granulocyte’, ‘leucocyte’, ‘immune cell’) or present in only one tissue (i.e. ‘erythroid lineage cell’; eye, ‘myeloid cell’; pancreas/prostate). The ‘erythrocyte’ and ‘erythroid progenitor’ annotations were further grouped to minimize multicollinearity.

Using the entire cell type space spanning all four organ compartments, either 30 observations (e.g. measured cells) were randomly sampled or the maximum number of available observations if less than 30 were subsampled, whichever was greater.

Cell type annotations were then reassigned based on the “broader” categories from hierarchical clustering (‘coarsegrain.py’). Raw count values from the Decont-X adjusted layer were used to minimize signal spread contamination impacting differentially expressed genes^13^.

This subsampled counts matrix was then passed to the ‘Create Signature Matrix’ analysis module at cibersortx.stanford.edu, with the following parameters:

- Disable quantile normalization = True
- Min. expression = 0.25
- Replicates = 5
- Sampling = 0.5
- Kappa = 999
- q-value = 0.01
- No. barcode genes = 3000 - 5000
- Filter non-hematopoietic genes = False

The resulting basis matrix was used in our nu-SVR deconvolution code, available on Github, under the name “tsp_v1_basisMatrix.txt”.

Abbreviations (left) of grouped cell types (right) in Figs. 1D as follows:

- gland cell: “acinar cell of salivary gland/myoepithelial cell”
- respiratory ciliated cell: “ciliated cell/lung ciliated cell”
- prostate epithelia: “club cell of prostate epithelium/hillock cell of prostate epithelium/hillock-club cell of prostate epithelium”
- salivary/bronchial secretory cell: “duct epithelial cell/serous cell of epithelium of bronchus”
- intestinal enterocyte: “enterocyte of epithelium of large intestine/enterocyte of epithelium of small intestine/intestinal crypt stem cell of large intestine/large intestine goblet cell/mature enterocyte/ paneth cell of epithelium of large intestine/small intestine goblet cell”
- erythrocyte/erythroid progenitor: “erythrocyte/erythroid progenitor”
- fibroblast/mesenchymal stem cell: “fibroblast/mesenchymal stem cell”
- intestinal secretory cell: “intestinal enteroendocrine cell/paneth cell of epithelium of small intestine/transit amplifying cell of small intestine”
- ionocyte/luminal epithelial cell of mammary gland: “ionocyte/luminal epithelial cell of mammary gland”
- secretory cell: “mucus secreting cell/secretory cell/tracheal goblet cell”
- pancreatic alpha/beta cell: “pancreatic alpha cell/pancreatic beta cell”
- respiratory secretory cell: “respiratory goblet cell/respiratory mucous cell/serous cell of epithelium of trachea”
- basal prostate cell: “basal cell of prostate epithelia”

### Nu-SVR deconvolution

We formulated the cell free transcriptome as a linear summation of the cell types from which it originates^4,48^. With this formulation, we adapted existing deconvolution methods developed with the objective of decomposing a bulk tissue sample into its single cell constituents^11,12^, where the deconvolution problem is formulated as:

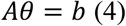

Here, *A* is the representative basis matrix (g x c) of **g** genes for **c** cell types, which represent the gene expression profiles of the **c** cell types. ***θ*** is a vector (c x 1) of the contributions of each of the cell types and **b** is the measured expression of the genes observed in from blood plasma (g x 1). The goal here is to learn ***θ*** such that the matrix product *Aθ* predicts the measured signal **b**. The derivation of the basis matrix *A* is described in the section ‘Basis Matrix Formation’.

We performed nu-SVR using a linear kernel to learn ***θ*** from a subset of genes from the signature matrix to best recapitulate the observed signal **b**, where nu denotes the lower bound on the fraction of support vectors and the upper bound on the fraction of errors at the margin^49^. Here, the support vectors are the genes used from the basis matrix from which to learn ***θ***; ***θ*** reflects the weights of the cell types in the basis matrix column space. For each sample, we learned coefficients for six values of nu, *ν* ∈ {0.05, 0.1, 0.15, 0.25, 0.5, 0.75} and five values of C, *C* ∈ {0.1, 0.5, 0.75, 1, 10} and estimated the resulting deconvolution error using the root mean square error (RMSE). We determined the product of the basis matrix with the learned coefficients (*Aθ*), which reflected some predicted expression value for each of the genes in a given cfRNA mixture used in the deconvolution. The RMSE was then computed using the predicted expression values and the measured values of these genes.

Only CPM counts ≥ 1 were considered in the mixture. The values in the basis matrix were also CPM-normalized. Prior to deconvolution, the mixture and basis matrix were scaled to zero mean and unit variance for improved runtime performance. We emphasize that we did not log-transform counts in *b* or in *A*, as this would destroy the requisite linearity assumption in equation 4. Specifically, the concavity of the log function would result in the consistent underestimation of ***θ*** during deconvolution^50^.

Using the ***θ*** resulting from the value of *ν* whose coefficients yielded the smallest RMSE was transformed to. Specifically, the relative fractional contributions of cell type specific RNA from ***θ***, we repeat what was previously described ^11,12^:

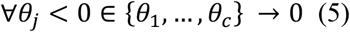

All non-zero coefficients were then normalized by their sum to result in the relative fractions to determine the relative fractional contributions of cell type specific RNA.

We used the function nuSVR from scikitlearn^51^ version 0.23.2.

The samples used for nu-SVR deconvolution were 75 non-cognitively impaired (NCI) patients from Toden et al spanning four sample collection centers. Given center-specific batch effects reported by Toden et al, we report our results on a per-center basis (Fig 1D, fig S5 & S6). There was good pairwise similarity of the learned coefficients amongst biological replicates within and across sample centers (fig. S6A, S6B). Deconvolution performance yielded RMSE and pearson r consistent with deconvolved GTEx tissues (fig. S3) whose distinct cell types were in the basis matrix column space (fig. S6C, S6D). In interpreting the resulting cell type fractions, a limitation of nu-SVR is that it uses highly expressed genes as support vectors, and consequently assigns a reduced fractional contribution to cell types expressing genes at lower levels or that are smaller in cell volume. Comparison of nu-SVR to quadratic programming^4^ and non-negative linear least squares^52^ yielded decreased deconvolution RMSE and comparable Pearson correlation. Furthermore, the cell type contributors identified with nu-SVR were consistent with the cell type markers detected using PanglaoDB in contrast to the other methods.

### Evaluating Basis Matrix on GTEx samples

Bulk RNA-seq samples from GTEx v8 were deconvolved with the derived basis matrix from tissues that were present (i.e. kidney cortex, whole blood, lung, spleen) or absent (e.g. kidney medulla and brain) from the basis matrix derived using Tabula Sapiens version 1. For each tissue type, the maximum number available samples or thirty samples, whichever was smaller, was deconvolved. Please see Supplementary Note 1 for additional discussion.

### Identifying tissue specific genes in cfRNA absent from basis matrix

To identify cell type specific genes in cfRNA that were distinct to a given tissue, we considered the set difference of the non-zero genes measured in a given cfRNA sample with the row space of the basis matrix and intersected this with HPA tissue specific genes:

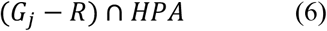

Where *G_j_* is the gene set in the j^th^ deconvolved sample, where a given gene in the set’s expression was ≥ 5 TMM-CPM. *R* is the set of genes in the row space of the basis matrix used for nu-SVR deconvolution. *HPA* denotes the total set of tissue specific genes from HPA.

The HPA tissue specific gene set (*HPA*) were genes across all tissues with Tissue Specificity assignments ‘Group Enriched’, ‘Tissue Enhanced’, ‘Tissue Enriched’ and NX expression ≥ 10. This approach yielded tissues with several distinct genes present in cfRNA which could then be subsequently interrogated using single cell data.

### Derivation of cell type specific gene profiles in context of the whole body using single cell data

For this analysis, only cell types unique to a given tissue (i.e. hepatocytes unique to the liver, or excitatory neurons unique to the brain) were considered so that bulk transcriptomic data could be used to ensure specificity in context of the whole body. A gene was asserted to be cell type specific if it was (i) differentially expressed within a given single cell tissue atlas (ii) possessed a Gini coefficient ≥ 0.6 and was listed as specific to the native tissue for the cell type of interest, indicating comprehensive tissue specificity in context of the whole body (fig. S8, S10).

#### (1) Single cell differential expression

For data received as a Seurat object, conversion to AnnData was performed by saving as an intermediate loom objects (Seurat version 3.1.5) and converting to AnnData (loompy version 3.0.6). Scanpy (version 1.6.0) was used for all other single cell analysis. Reads per cell were normalized for library size (scanpy normalize_total, target_sum = 1e4), then logged (scanpy log1p). Differential expression was performed using the Wilcoxon rank sum test in Scanpy’s filter_rank_genes_groups with the following arguments: min_fold_change = 1.5, min_in_group_fraction = 0.2, max_out_group_fraction = 0.5, corr_method = “benjamini-hochberg”. For differentially expressed genes (DEG) with Benjamini Hochberg adjusted p-values < 0.01, the ratio of the highest out_group percent expressed to in-group percent expressed < 0.5 to ensure high specific expression in the cell type of interest within a given cell type atlas.

#### (2) Quantifying comprehensive whole-body tissue specificity using the Gini coefficient

The distribution of all the Gini coefficients and Tau values across all genes belonging to cell type gene profiles for cell types native to a given tissue were compared using the HPA gene expression Tissue Specificity and Tissue Distribution assignments^15^ (fig. S9). The Gini coefficient better reflected the underlying distribution of gene expression tissue-specificity than Tau (fig. S9) and was hence used for subsequent analysis. As the Gini coefficient approaches unity, this indicates extreme gene expression inequality, or equivalently high specificity. A single threshold (Gini coefficient ≥ 0.6) was applied across all atlases to facilitate a generalizable framework from which to define tissue specific cell type gene profiles in context of the whole body in a principled fashion for signature scoring in cfRNA.

For the following definitions, *n* denotes the total number of tissues and *x_i_* is the expression of a given gene in the i^th^ tissue.

The Gini coefficient was computed as defined in ^53^:

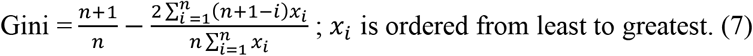

Tau, as defined in ^53^:

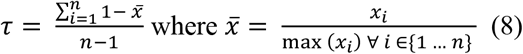

HPA NX Counts from the HPA object entitled ‘rna_tissue_consensus.tsv’ accessed on July 1, 2019 were used for computing Gini coefficients and Tau.

<u>Note for brain cell type gene profiles:</u> given that there are multiple sub brain-regions in the HPA data, the determined Gini coefficients are lower (e.g. not as close to unity compared to other cell type gene profiles) since there are multiple regions of the brain with high expression, which would result in reduced count inequality.

### Gene Expression in GTEx

We confirmed the specificity of a given gene profile to its corresponding cell type by comparing the aggregate expression of a given cell type signature in its native tissue compared to that of the average across remaining GTEx tissues (fig. S8C and fig. S10F, 10G). We uniformly observed a median fold change greater than one in the signature score of a cell type gene profile in its native tissue relative to the mean expression in other tissues, confirming high specificity.

Raw GTEx data v8 (accessed August 26 2019) was converted to log(counts-per-ten-thousand + 1) counts. The signature score was determined by summing the expression of the genes in a given bulk RNA sample for a given cell type gene profile. Since only gene profiles were derived for cell types that correspond to a given tissue, the mean signature score of a cell type profile across the non-native tissues was then computed and used to determine the log fold change.

### Cell Type specificity of differentially expressed genes in AD and NAFLD cfRNA

After observing a significant intersection between the DEG in AD/NCI^6^ or NAFLD^7^ in cfRNA with corresponding cell type specific genes (fig. S13A, S14A), we then assessed the cell type specificity of DEG using a permutation test. To assess whether DEGs that intersected with a cell type gene profile were more specific to a given cell type than DEGs that were generally tissue specific, we performed a permutation test. Specifically, we compared the Gini coefficient for genes in these two groups, computed using the mean expression of a given gene across brain cell types from healthy brain^28^ or liver^22^ single cell data. We considered the cell type gene profiles as defined for signature scoring in Fig. 2.

The starting set of tissue specific genes were defined using in the HPA tissue transcriptional data annotated as either ‘Tissue enriched’, ‘Group enriched’, or ‘Tissue enhanced’ (brain, accessed January 13, 2021; liver, accessed November 28, 2020). These requirements ensured the specificity of a given brain/liver gene in context of the whole body. This formed the initial set of tissue specific genes *B*.

The union of all brain cell type specific genes is the set *C*. All genes in *C* (‘cell type specific’) were a subset of the respective initial set of tissue specific genes:

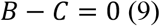

Genes in *B* that that did not intersect with *C* (e.g. any cell type gene profile (‘bcell type specific’)) and intersected with DEG-up (*U*) or DEG-down genes (*D*)^6^ were then defined as ‘tissue specific’.

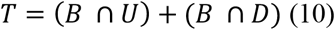

The Gini coefficients reflecting the gene expression inequality across the cell types within corresponding tissue single cell atlas were computed for the gene sets labelled as ‘cell type specific’ and ‘tissue specific’. Brain reference data to compute Gini coefficients was the single cell brain atlas with diagnosis as ‘Normal’^28^. Liver tissue data was used as-is. All Gini coefficients were computed using the mean log transformed CPTT (counts per ten thousand) gene expression per cell type.

A permutation test was then performed on the union of the Gini coefficients for the genes labeled as ‘brain cell type specific’ and ‘brain tissue specific’. The purpose of this test was to assess probability that the observed mean difference in Gini coefficient for these two groups yielded no difference in specificity (e.g. H_0_: *μ_cell type Gini Coefficient_* = *μ_tissue Gini coefficient_*).

Gini coefficients were permuted and reassigned to the list of ‘brain tissue’ or ‘brain cell type’ genes, then the difference in mean of the two groups was computed. This procedure was repeated 10,000 times. The p-value was determined as follows:

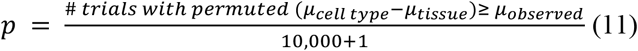

Where *μ_observed_* ≔ (*μ_cell type Gini Coefficient_* − *μ_brain tissue Gini coefficient_*).

The additional 1 in the denominator reflects the original test between the true difference in means (e.g. the true comparison yielding *μ_observed_*)

NAFLD: We considered the space of reported NAFLD differentially expressed genes in serum^7^. Here *C* = hepatocyte gene profile and *B* = the liver specific genes.

AD: First, we intersected a given cell type gene profile in AD with the equivalent NCI profile for comparative analysis. Genes defined as ‘brain cell type specific’ for signature scoring in Fig. 2D were used in this comparison. Of note, no DEG-up genes intersected with any of the brain cell type signatures in Fig 2D. Microglia, though often implicated in AD pathogenesis were excluded given their high overlapping transcriptional profile with non-central nervous system macrophages^54^. Inhibitory neurons were also excluded given the low number of cell type specific genes intersecting between AD and NCI phenotypes.

### Estimating signature scores for each cell type

The signature score is defined as the sum of the log-transformed CPM-TMM normalized counts per gene asserted to be cell type specific, where *i* denotes the index of the gene in a cell type signature gene profile in the *j*^th^ patient sample:

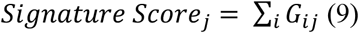

<u>Preeclampsia:</u> for signature scoring of syncytiotrophoblast and extravillous trophoblast gene profiles in PEARL-PEC and iPEC^5^, a respective cell type gene profile used for signature scoring was derived as described in ‘Derivation of cell type specific gene profiles in context of the whole body using single cell data’ independently using two different placental single cell datasets^19,20^. Only the intersection of the cell type specific gene profiles for a given trophoblast cell type between the two datasets was included in the respective trophoblast gene profile for signature scoring.

#### Chronic Kidney Disease

We compared the signature score of the proximal tubule in CKD (9 patients; 51 samples) and healthy controls (3 patients; 9 samples). Given that all patients samples were longitudinally sampled over ~30 days (e.g. individual samples were taken on different days), we treated the samples as biological replicates and included all time points since the timescale over which renal cell type changes typically occur is longer than the collection period. The sequencing depth was comparable between the CKD and healthy cohorts though it was reduced in comparison to the other cfRNA datasets used in this work. To account for gene measurement dropout, we required that the expression of a given gene in the proximal tubule gene profile was non-zero in at least one sample in both cohorts. Given that all samples were sequenced together, no batch correction was necessary, facilitating a representative comparison between CKD and healthy.

#### Alzheimer’s Disease

Microglia, though often implicated in AD pathogenesis were excluded given their high overlapping transcriptional profile with non-central nervous system macrophages^54^. Inhibitory neurons were also excluded given the low number of cell type specific genes intersecting between AD and NCI phenotypes. Brain gene profiles as defined in the AD section of ‘*Cell Type specificity of differentially expressed genes in AD and NAFLD cfRNA’* were utilized.

### Assessing p-value calibration for a given signature score

Cell type signature scores were tested between control and sick samples with a Mann-Whitney U test. The resulting p-values were calibrated with a permutation test. Here, the labels compared in a given test (i.e. CKD vs. CTRL, or AD vs. NCI, NAFLD vs. CTRL, etc.) were randomly shuffled 10,000 times. We observed a well-calibrated, uniform p-value distribution (fig. S12–S14), validating the experimentally observed test statistics.

## Supplementary Note 1: Deconvolution of bulk GTEx tissues using the *Tabula Sapiens*-derived basis matrix

To assess the ability of the basis matrix to deconvolve tissues whose cell types were wholly present in the cell type column space, we deconvolved a subset of bulk RNA-seq GTEx samples. The determined fractions of cell type specific RNA recapitulated the predominant cell types within a given tissue (fig. S4). Organs with increased cell type heterogeneity (lung, bladder, kidney, intestine, colon) in contrast to tissues with reduced spatial heterogeneity (liver, spleen, whole blood)^13^, exhibited greater variance in deconvolved fractions (fig. S4) and deconvolution performance (fig. S3). Tissues with reduced spatial heterogeneity whose cell types were wholly in the basis matrix column space include predominantly b cells/plasma cells and erythrocytes in spleen (fig. S4F); hepatocytes, liver (fig. S4C); erythrocytes and leukocytes, whole blood (fig. S4D). Cell types belonging to tissues with increased spatial heterogeneity exhibited greater variance in deconvolved fractions: kidney cortex majority fractions were from kidney epithelia and lymphocytes (fig. 4G); small intestine, intestinal enterocytes and lymphocytes (fig. S4I); lung, pneumonocytes and immune cells (fig. S4J), colon, intestinal enterocytes, lymphocytes, and muscle cells (fig. S4E). Cells with larger volume yielded larger deconvolved fractions across all tissues (fig. S4). Variance in the relative cell type fractional contributions across the deconvolved bulk samples within a given tissue reflects the underlying cell type heterogeneity, particularly in these complex samples. GTEx kidney medulla samples recorded to be contaminated with renal cortex reflect the presence of the kidney epithelia, the majority cell type in the renal cortex (fig. S4G). Given that the kidney medulla is not part of TSP v1.0, we did not expect high deconvolution performance since its cell types are absent from the basis matrix column space. The brain, whose cell types were wholly absent from the cell type column space exhibited poor deconvolution performance, as expected (fig. S3, S4B). However, the majority cell type fraction assigned was to the cell type belonging to the peripheral nervous system, the schwann cell, underscoring the ability of our deconvolution method to assign fractional contributions to similar cell types from those that are absent from the basis matrix column space.

## Supplementary Note 2: Noninvasive measurement of trophoblast cell type signatures in preeclampsia

In pregnancy, extravillous trophoblast (EVT) invasion is a stage in uteroplacental arterial remodeling^20,23^. Arterial remodeling occurs to ensure adequate maternal blood flow to the growing fetus^20,23^ and is sometimes reduced in preeclampsia^23^. Previously, EVT was reported by Tsang et al to be noninvasively resolvable and elevated in early onset preeclampsia (gestational age at diagnosis < 34 weeks) as compared to healthy pregnancy^24^. However, examination of the trophoblast gene profiles used by Tsang et al. using two independent placental single-cell atlases^19,20^ revealed several genes that were not cell type specific or exhibited very low trophoblast expression (fig. S11C, 11D), thereby adversely impacting signature score interpretation.

*CERCAM*, *IL18BP*, and *PYCR1* are not extravillous trophoblast specific, exhibiting higher expression in fibroblast cell types in both atlases, despite Tsang’s inclusion in their EVT gene profile (fig. S11C, 11D). Furthermore, EVT genes in Tsang’s gene profile, *RRAD*, *SLC6A2*, and *UPK1B* all exhibit very low EVT expression across both placental atlases. Numerous PSG genes (*PSG11*, *PSG1/PSG2*, *PSG3*, *PSG4*, *PSG6*, *PSG9*) do not exhibit high syncytiotrophoblast (SCT) expression, despite their inclusion in Tsang’s SCT gene profile. GH2 either exhibits no expression or comparable non-SCT specific expression across cell types in both atlases (fig. S11C, 11D).

The presence of these non-cell type specific genes in a cell type gene profile consequently impacted the interpretation of Tsang et al’s signature scores. Using our criteria for deriving a given cell type gene profile (Methods), we derived gene profiles for the same two cell types, EVT and SCT (fig. S10), and then quantified their respective signature scores in two previously published preeclampsia cohorts^5^ (fig. S11). In contrast to Tsang et al, we observed no significant difference in either trophoblast signature score in cfRNA samples collected at diagnosis for mothers with early-onset preeclampsia (fig. S11) (p = 0.703 and U = 1396, 0.794 and U = 1416 respectively, two-sided Mann Whitney U) and for mothers with either early- or late-onset preeclampsia (p = 0.24 and H = 4.18, 0.54 and H =2.15 respectively, Kruskal Wallace, fig. S10B) as compared to samples from mothers with no complications at a matched gestational age.

In our work deriving cell type gene profiles for signature scoring in cfRNA, we only considered genes with high log fold change in a given cell type population and low expression in any other measured cell type (Methods). We acknowledge that this method may miss some genes for a given cell type population with low uniform expression (i.e. low expression in a large fraction of cells of a given type) or with heterogeneous expression (i.e. high expression in a small fraction of cells of a given type). However, since this work is the first comprehensive examination of cell type specific origins in the cell free transcriptome, we sought to be conservative in what we asserted to be cell type specific so that we could be confident in measuring a cell type signature score noninvasively; this approach boded well for all diseases presented in our work.

Taken together with validation in two independent placental cell atlases, we conclude that the EVT and SCT cell type gene profiles by Tsang et al. do not enable estimation of trophoblast pathology from cfRNA in preeclampsia. The role of extravillous trophoblast invasion and the ubiquity of its cellular pathophysiology in preeclampsia thus remains an open question.

## Tabula Sapiens Consortium

### Overall Project Direction and Coordination

Robert C Jones, Jim Karkanias, Mark Krasnow, Angela Oliveira Pisco, Stephen Quake, Julia Salzman, Nir Yosef

### Donor Recruitment

Bryan Bulthaup, Phillip Brown, Will Harper, Marisa Hemenez, Ravikumar Ponnusamy, Ahmad Salehi, Bhavani Sanagavarapu, Eileen Spallino

### Surgeons

Ksenia A. Aaron, Waldo Concepcion, James Gardner, Burnett Kelly, Nikole Neidlinger, Zifa Wang

### Logistical coordination

Sheela Crasta, Saroja Kolluru, Maurizio Morri, Angela Oliveira Pisco, Serena Y. Tan, Kyle J. Travaglini, Chenling Xu

### Organ Processing

Marcela Alcántara-Hernández, Nicole Almanzar, Jane Antony, Benjamin Beyersdorf, Deviana Burhan, Lauren Byrnes, Kruti Calcuttawala, Mathew Carter, Charles K. F. Chan, Charles A. Chang, Alex Colville, Sheela Crasta, Rebecca Culver, Ivana Cvijović, Jessica D’Addabbo, Gaetano D’Amato, Camille Ezran, Francisco Galdos, Astrid Gillich, William R. Goodyer, Yan Hang, Alyssa Hayashi, Sahar Houshdaran, Xianxi Huang, Juan Irwin, SoRi Jang, Julia Vallve Juanico, Aaron M. Kershner, Soochi Kim, Bernhard Kiss, Saroja Kolluru, William Kong, Maya Kumar, Rebecca Leylek, Baoxiang Li, Shixuan Liu, Gabriel Loeb, Wan-Jin Lu, Shruti Mantri, Maxim Markovic, Patrick L. McAlpine, Ross Metzger, Antoine de Morree, Maurizio Morri, Karim Mrouj, Shravani Mukherjee, Tyler Muser, Patrick Neuhöfer, Thi Nguyen, Kimberly Perez, Ragini Phansalkar, Angela Oliveira Pisco, Nazan Puluca, Zhen Qi, Poorvi Rao, Hayley Raquer, Koki Sasagawa, Nicholas Schaum, Bronwyn Lane Scott, Bobak Seddighzadeh, Joe Segal, Sushmita Sen, Sean Spencer, Lea Steffes, Varun R. Subramaniam, Aditi Swarup, Michael Swift, Kyle J Travaglini, Will Van Treuren, Emily Trimm, Maggie Tsui, Sivakamasundari Vijayakumar, Kim Chi Vo, Sevahn K. Vorperian, Hannah Weinstein, Juliane Winkler, Timothy T.H. Wu, Jamie Xie, Andrea R.Yung, Yue Zhang

### Sequencing

Angela M. Detweiler, Honey Mekonen, Norma Neff, Rene V. Sit, Michelle Tan, Jia Yan

### Histology

Gregory R. Bean, Gerald J. Berry, Vivek Charu, Erna Forgó, Brock A. Martin, Michael G. Ozawa, Oscar Silva, Serena Y. Tan, Pranathi Vemuri

### Computational Data Analysis

Shaked Afik, Rob Bierman, Olga Botvinnik, Ashley Byrne, Michelle Chen, Roozbeh Dehghannasiri, Angela Detweiler, Adam Gayoso, Qiqing Li, Gita Mahmoudabadi, Aaron McGeever, Antoine de Morree, Julia Olivieri, Madeline Park, Angela Oliveira Pisco, Neha Ravikumar, Julia Salzman, Geoff Stanley, Michael Swift, Michelle Tan, Weilun Tan, Sevahn K. Vorperian, Sheng Wang, Galen Xing, Chenling Xu, Nir Yosef

### Expert Cell Type Annotation

Marcela Alcántara-Hernández, Jane Antony, Charles A. Chang, Alex Colville, Sheela Crasta, Rebecca Culver, Camille Ezran, Astrid Gillich, Yan Hang, Juan Irwin, SoRi Jang, Aaron M. Kershner, William Kong, Rebecca Leylek, Gabriel Loeb, Ross Metzger, Antoine de Morree, Patrick Neuhöfer, Kimberly Perez, Ragini Phansalkar, Zhen Qi, Hayley Raquer, Bronwyn Lane Scott, Rahul Sinha, Hanbing Song, Sean Spencer, Aditi Swarup, Michael Swift, Kyle J. Travaglini, Jamie Xie

### Tissue Expert Principal Investigators

Steven E. Artandi, Philip Beachy, Michael F. Clarke, Linda Giudice, Franklin Huang, KC Huang, Juliana Idoyaga, Seung K Kim, Mark Krasnow, Christin Kuo, Patricia Nguyen, Stephen Quake, Thomas A. Rando, Kristy Red-Horse, Jeremy Reiter, Justin Sonnenburg, Bruce Wang, Albert Wu, Sean Wu, Tony Wyss-Coray

---

**Fig. S1.**
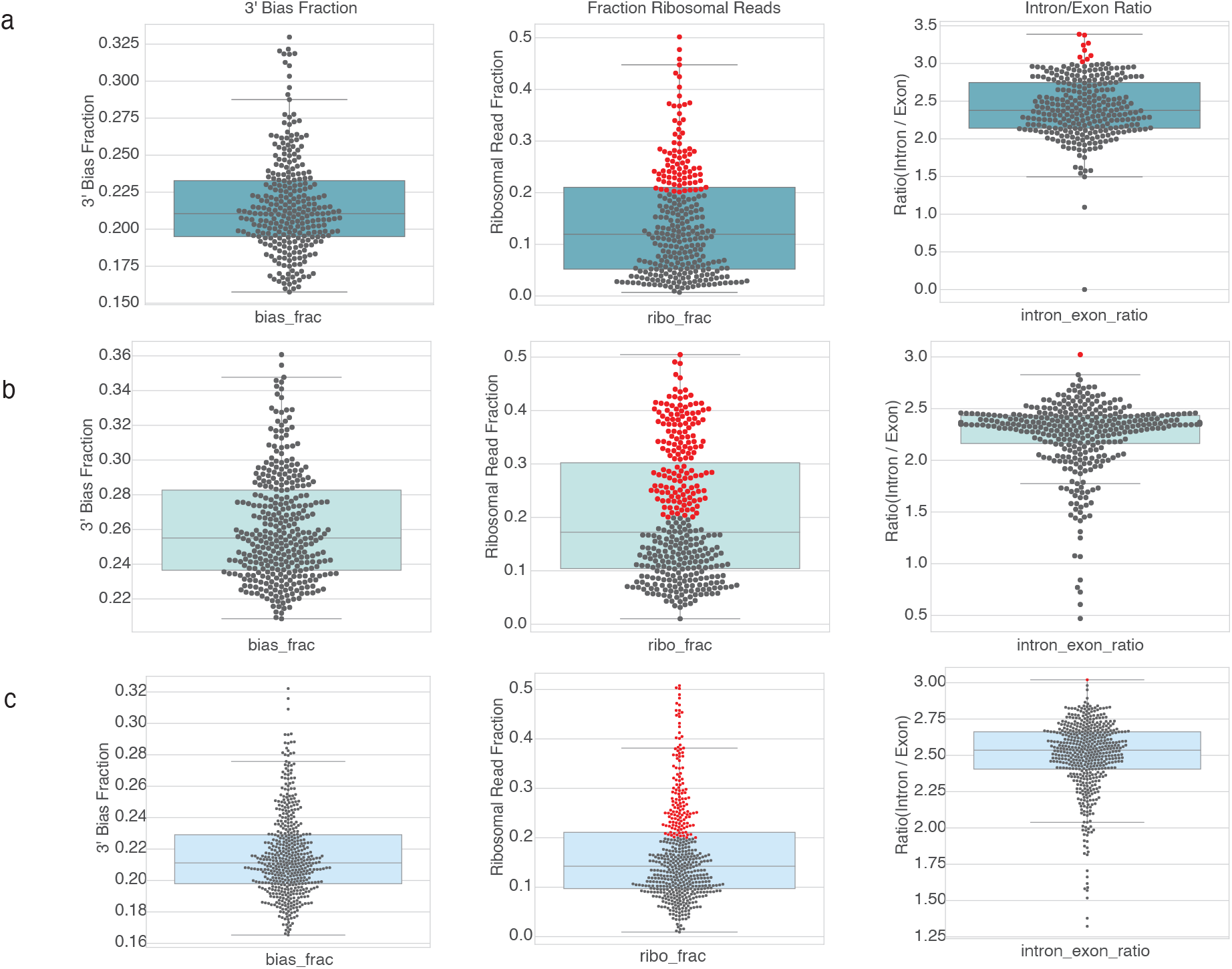
Identification of samples with outlier values for at least one quality control metric including a measure of RNA degradation, ribosomal fraction, and DNA contamination from Ibarra et al. Samples with outlier values are highlighted in red. (See Methods section ‘Sample quality filtering’ for details). Box plot: horizonal line, median; lower hinge, 25^th^ percentile; upper hinge, 75^th^ percentile; whiskers span the 1.5 interquartile range; points outside the whiskers indicate outliers. Panels represent all published cfRNA samples from (a) Ibarra et al (b) Toden et al (c) Chalasani et al.

**Fig. S2.**
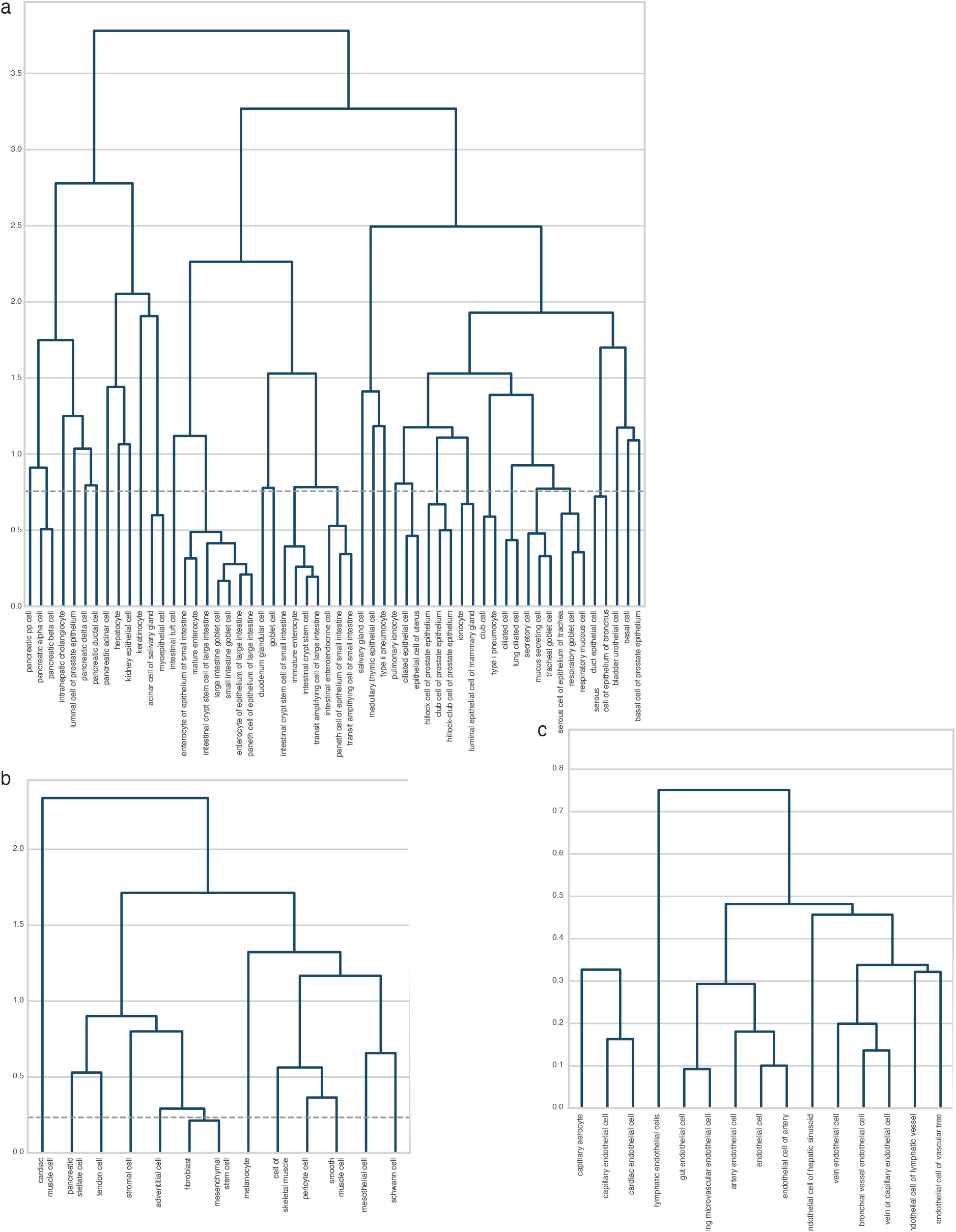
Hierarchical clustering on non-immune Tabula Sapiens organ compartments for annotation coarsegraining in basis matrix generation. Dashed line indicates height at which tree was cut. Dendrograms correspond with the cell type annotations belonging to the (a) epithelial compartment, (b) the stromal compartment, (c) the endothelial compartment.

**Fig. S3.**
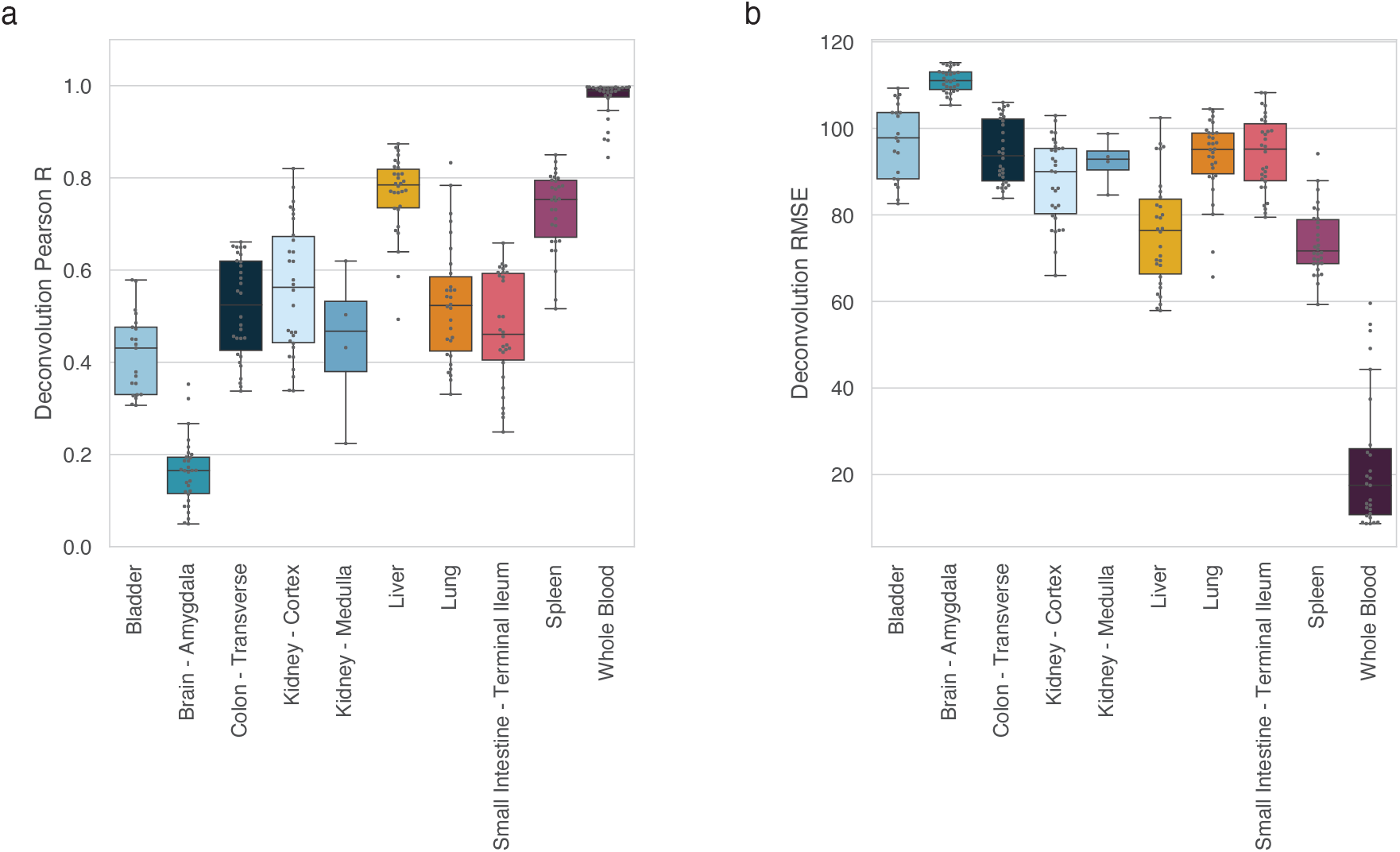
Basis matrix performance on GTEx bulk RNA samples using nu-SVR. GTEx tissue samples possessing cell types wholly present (Kidney – Cortex, Spleen, Small Intestine – Terminal Ileum, Lung, Whole Blood) and absent from the basis matrix column space (Kidney – Medulla, Liver) were selected. For box plots: horizonal line, median; lower hinge, 25^th^ percentile; upper hinge, 75^th^ percentile; whiskers,1.5 interquartile range; points outside the whiskers indicate outliers. (a) Pearson correlation between predicted expression and actual expression in cfRNA. (b) Root Mean Square Error between predicted expression and actual expression in cfRNA. Units are zero-mean unit variance scaled CPM counts; tissues present in TSP have reduced RMSE compared to those that are absent (e.g. Kidney – Medulla and Brain). Tissues with high cellular heterogeneity (Lung, Bladder, Small Intestine, Kidney) exhibit reduced deconvolution performance compared to simpler tissues (Whole Blood, Spleen, Liver)

**Fig S4.**
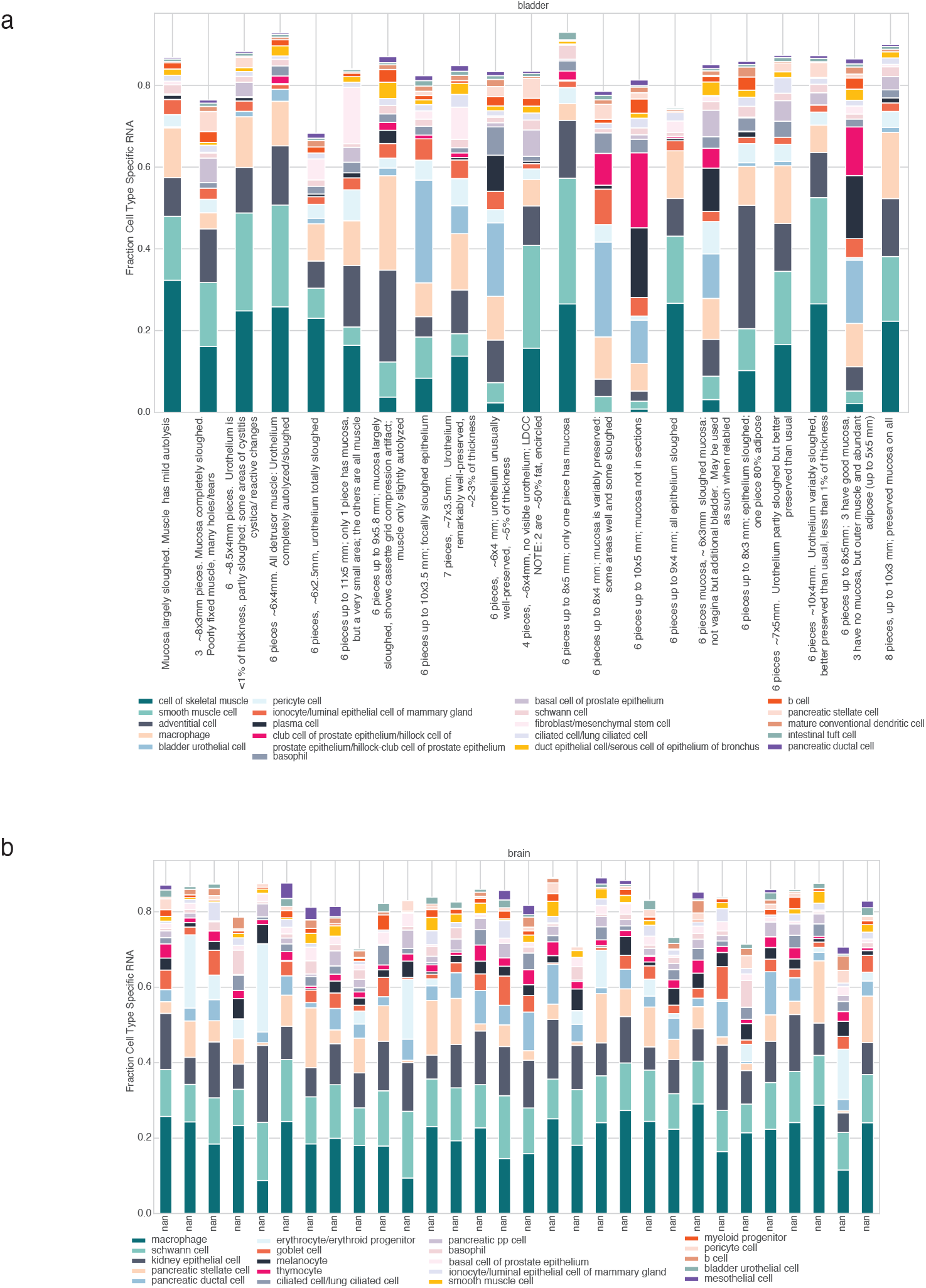
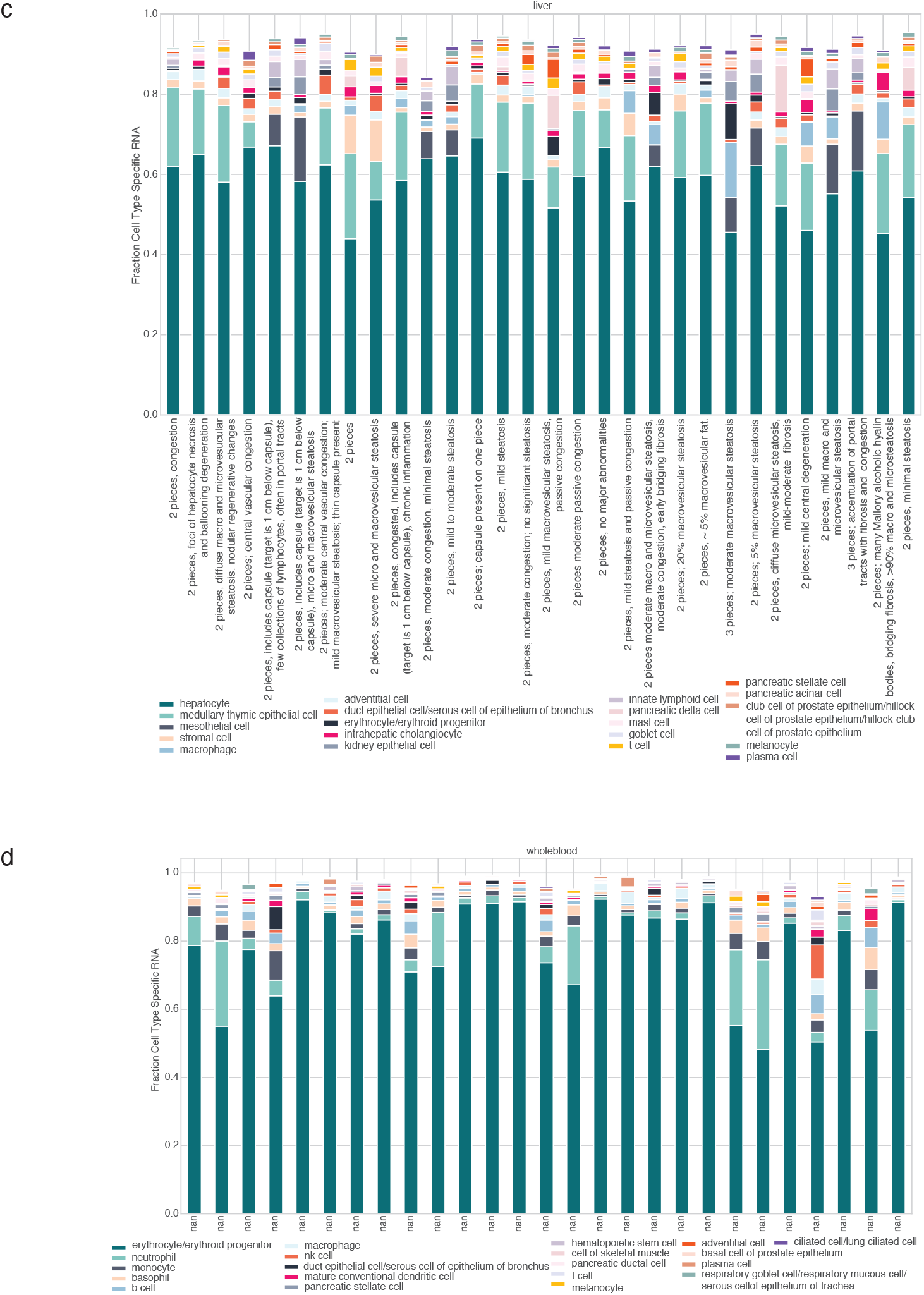
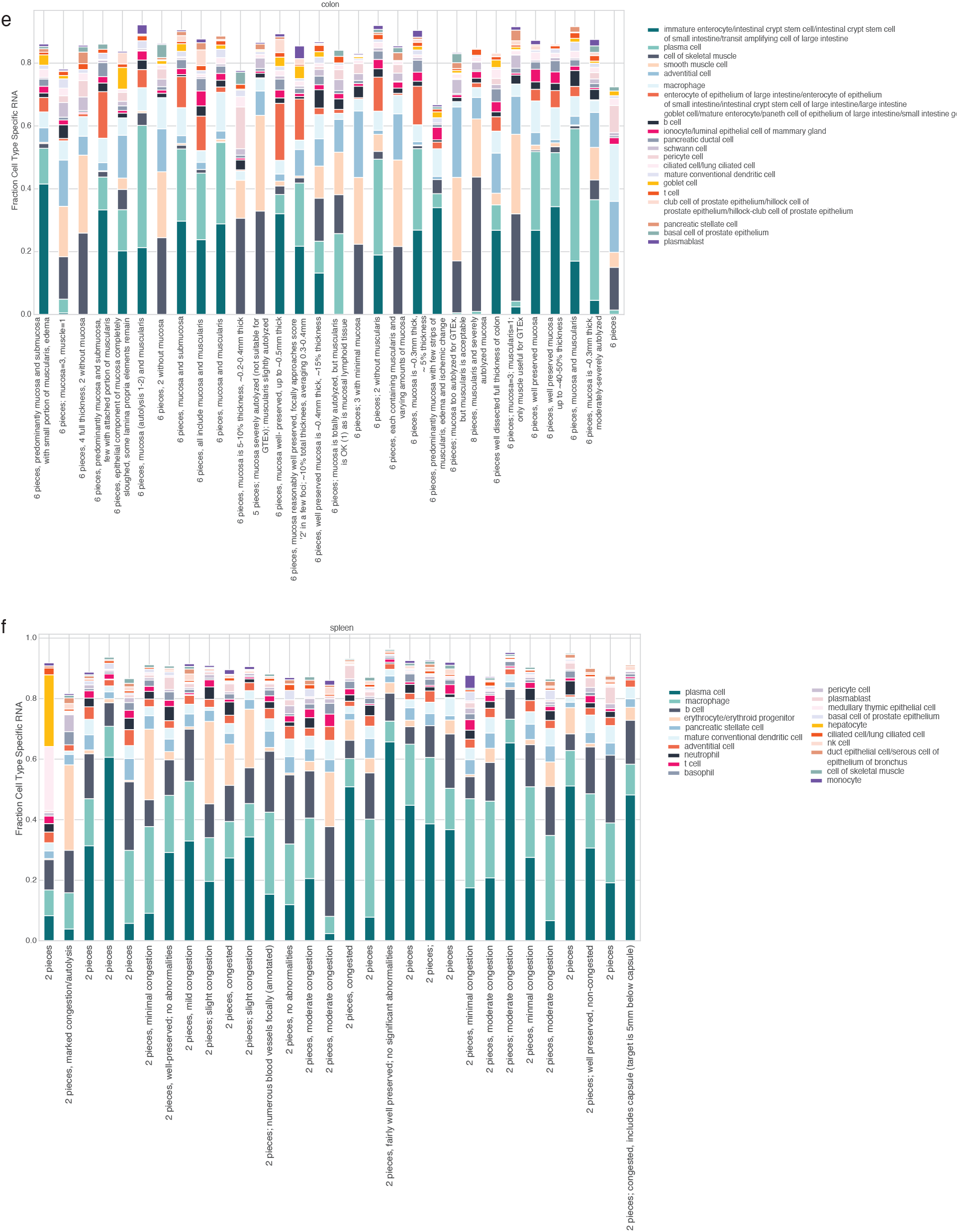
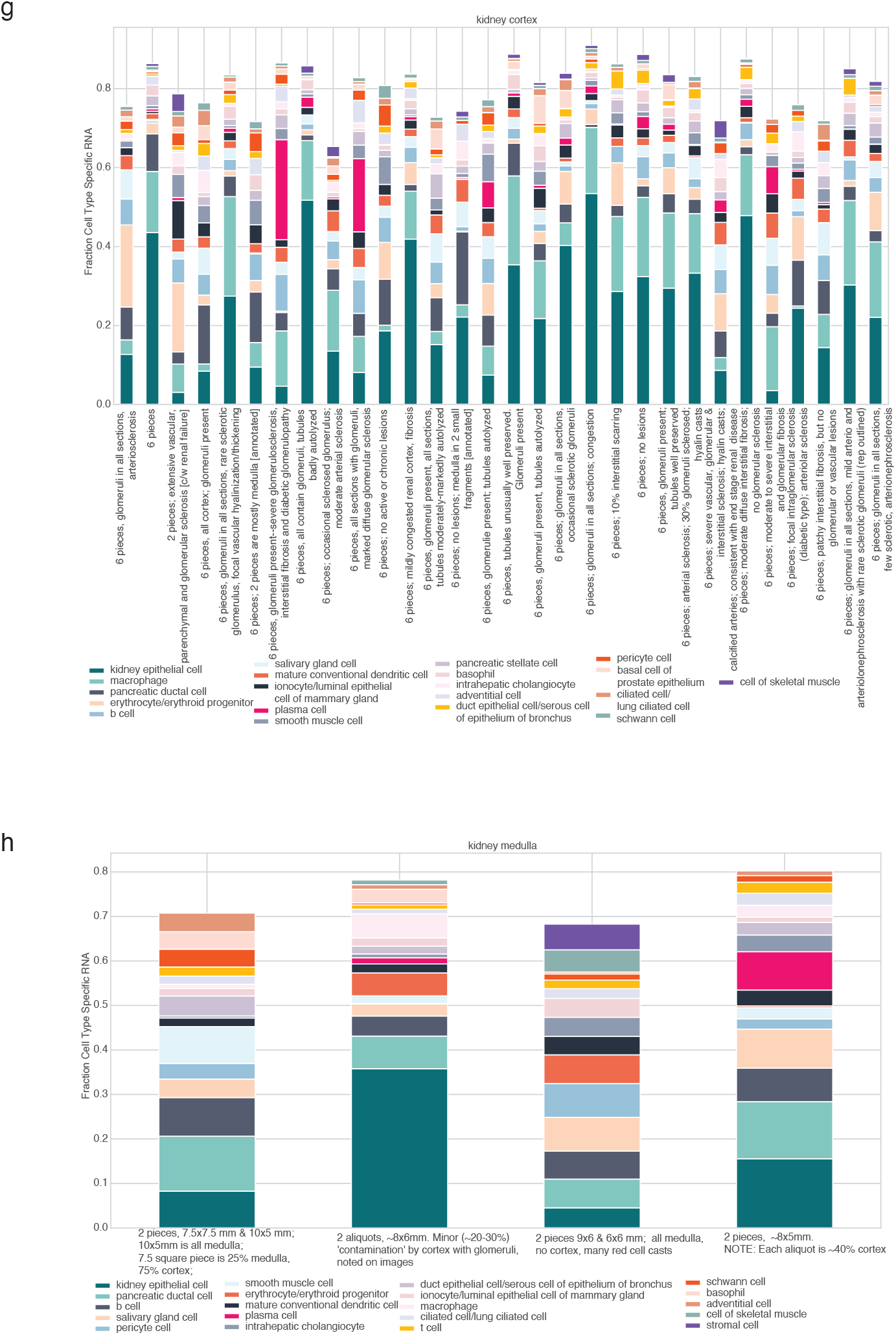
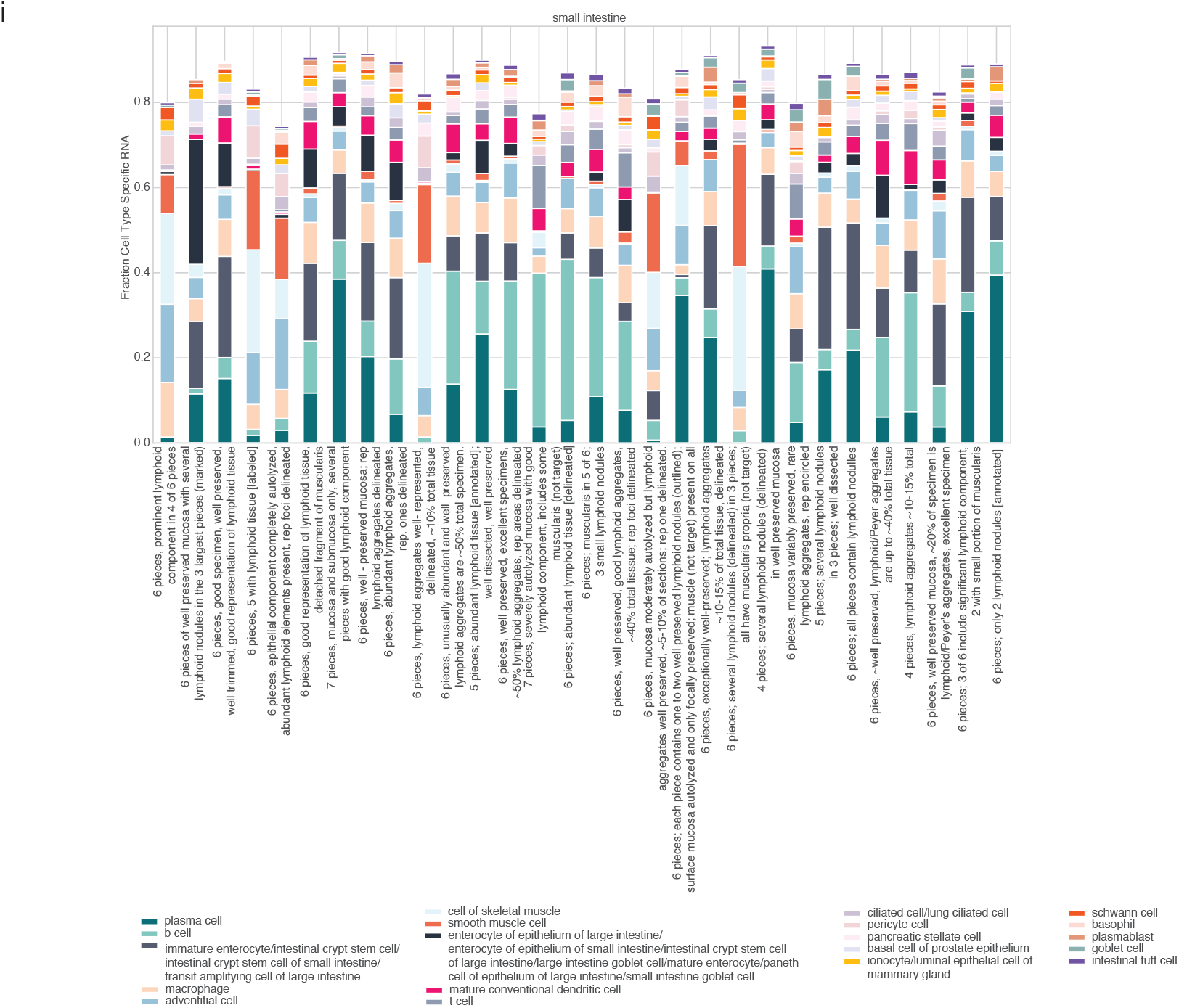
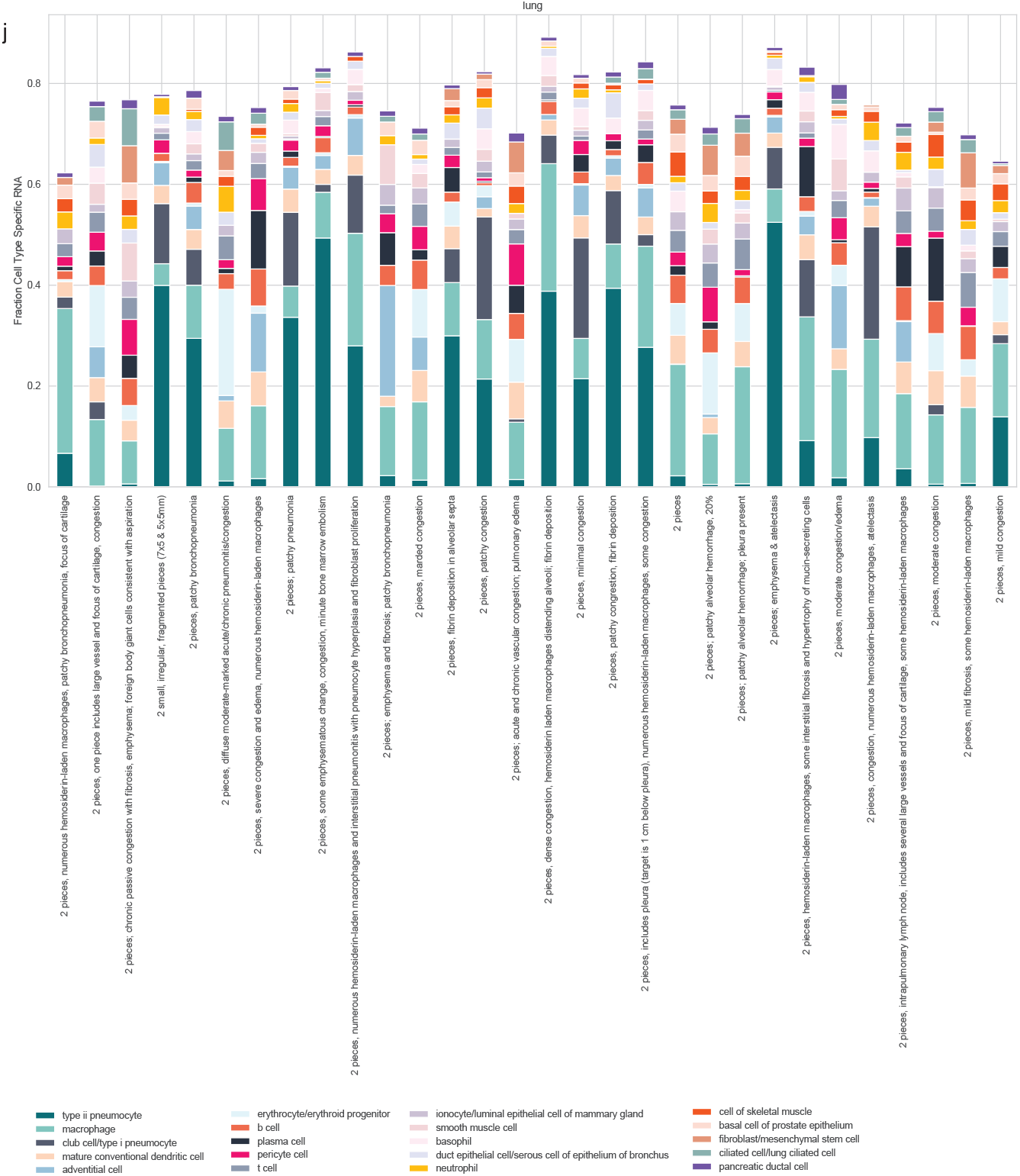
Deconvolved fractions of cell type specific RNA from various GTEx tissues using nu-SVR to assess deconvolution performance of the *Tabula Sapiens*-derived basis matrix. The two tissues whose cell types were absent from the basis matrix column space were Kidney – Medulla and Brain. Kidney medulla samples reported to be contaminated with cortex are reflected by deconvolved kidney epithelia fractions. The brain, which is absent from the TSP v1.0, has majority fractions of schwann cells, a peripheral nervous cell type. Majority cell types for a given tissue, such as lung pneumonocytes and immune cells in the lung or kidney epithelia for the kidney cortex underscore the ability for the signature matrix to capture representative fractions of cell type specific RNA and reflect underlying cell heterogeneity in bulk RNA-seq data. Additional discussion is in Supplementary Note 1. (a) Bladder (b) Brain (c) Liver (d) Whole Blood (e) Colon – Transverse (f) Spleen (g) Kidney – Cortex (h) Kidney – Medulla (i) Small Intestine – Terminal Ileum (j) Lung

**Fig. S5.**
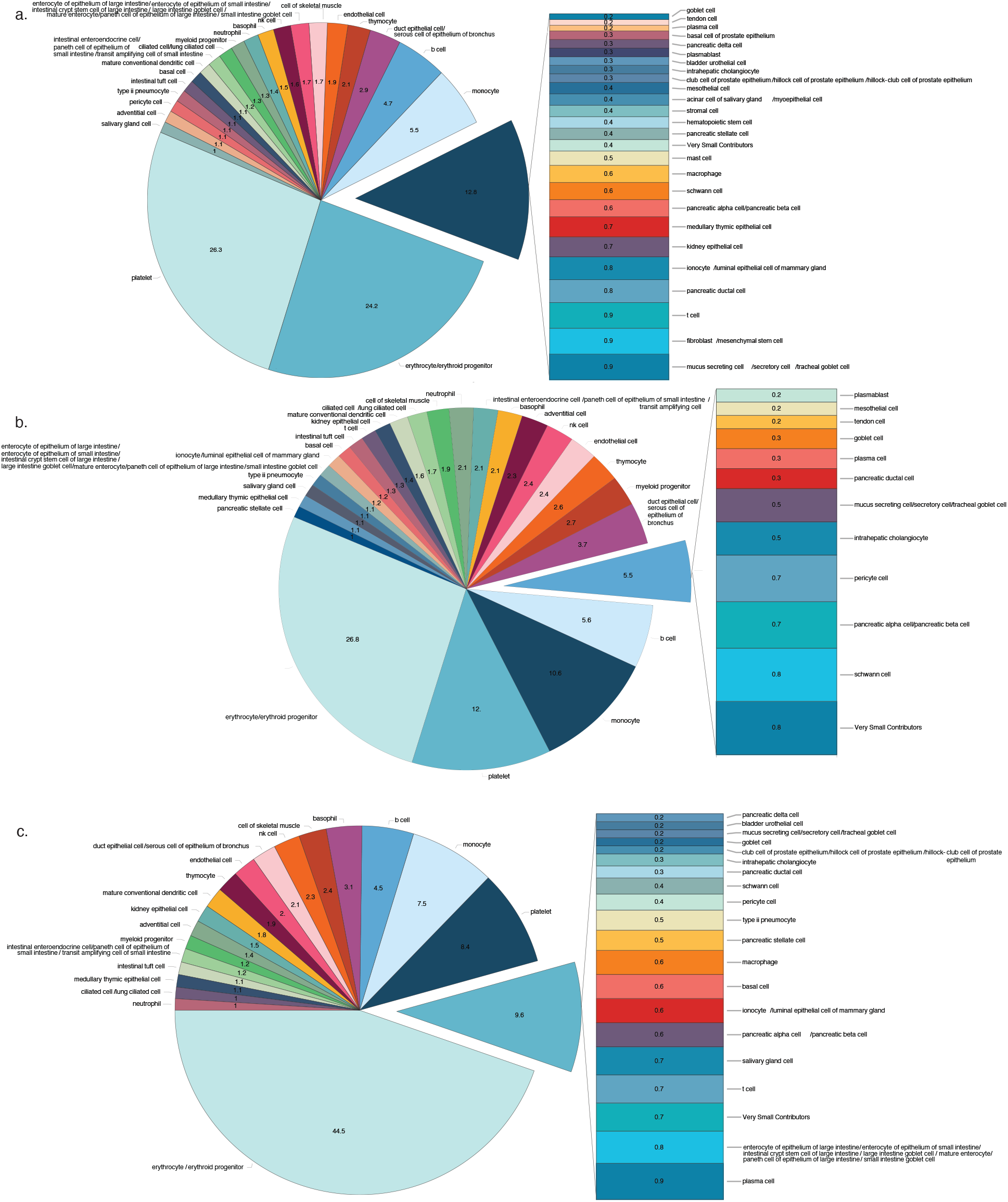
Deconvolution of healthy plasma samples from Toden et al using Tabula Sapiens. Pie charts denote mean fractional cell type specific RNA contributions for (a) BioIVT (n = 18), (b) University of Indiana (n = 17), (c) Washington University in St. Louis (n = 22).

**Fig. S6.**
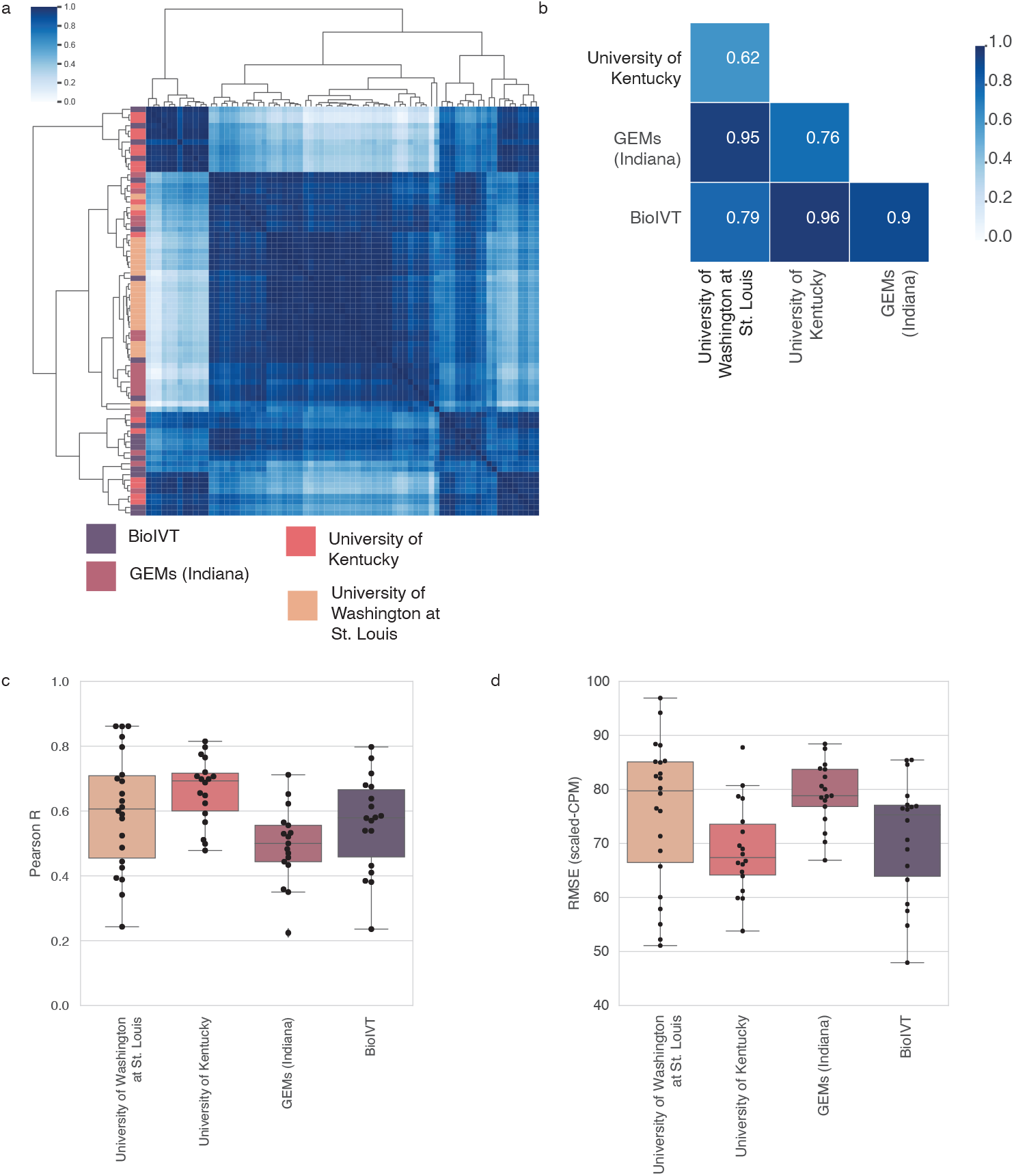
Deconvolving the plasma cell free transcriptome with Tabula Sapiens using nu-SVR (n = 75 samples). For boxplots, Horizonal line, median; lower hinge, 25^th^ percentile; upper hinge, 75^th^ percentile; whiskers span the 1.5 interquartile range; points outside the whiskers indicate outliers. (a) Complete linkage clustermap of pairwise pearson correlation of deconvolved cell type fractions between patients from a given center; row color denotes a given center. (b) Heatmap of pairwise pearson correlation of the mean cell type coefficients per center (c) Deconvolution pearson correlation between predicted vs. measured expression for all biological replicates across all centers. (d) Deconvolution RMSE between predicted vs. measured expression for all biological replicates across all centers.

**Fig S7.**
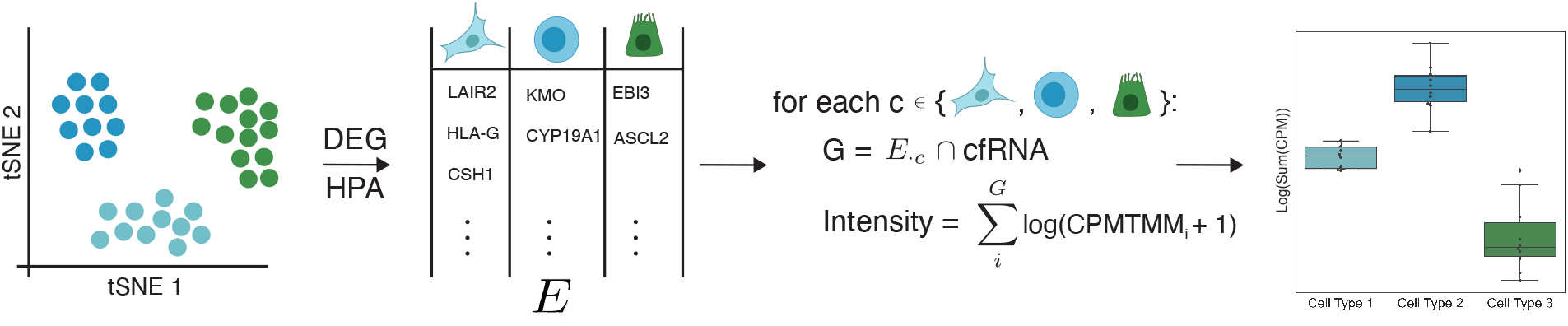
Cell type signature scoring procedure. See ‘Signature Scoring’ section of methods for derivation of cell type gene profiles.

**Fig S8.**
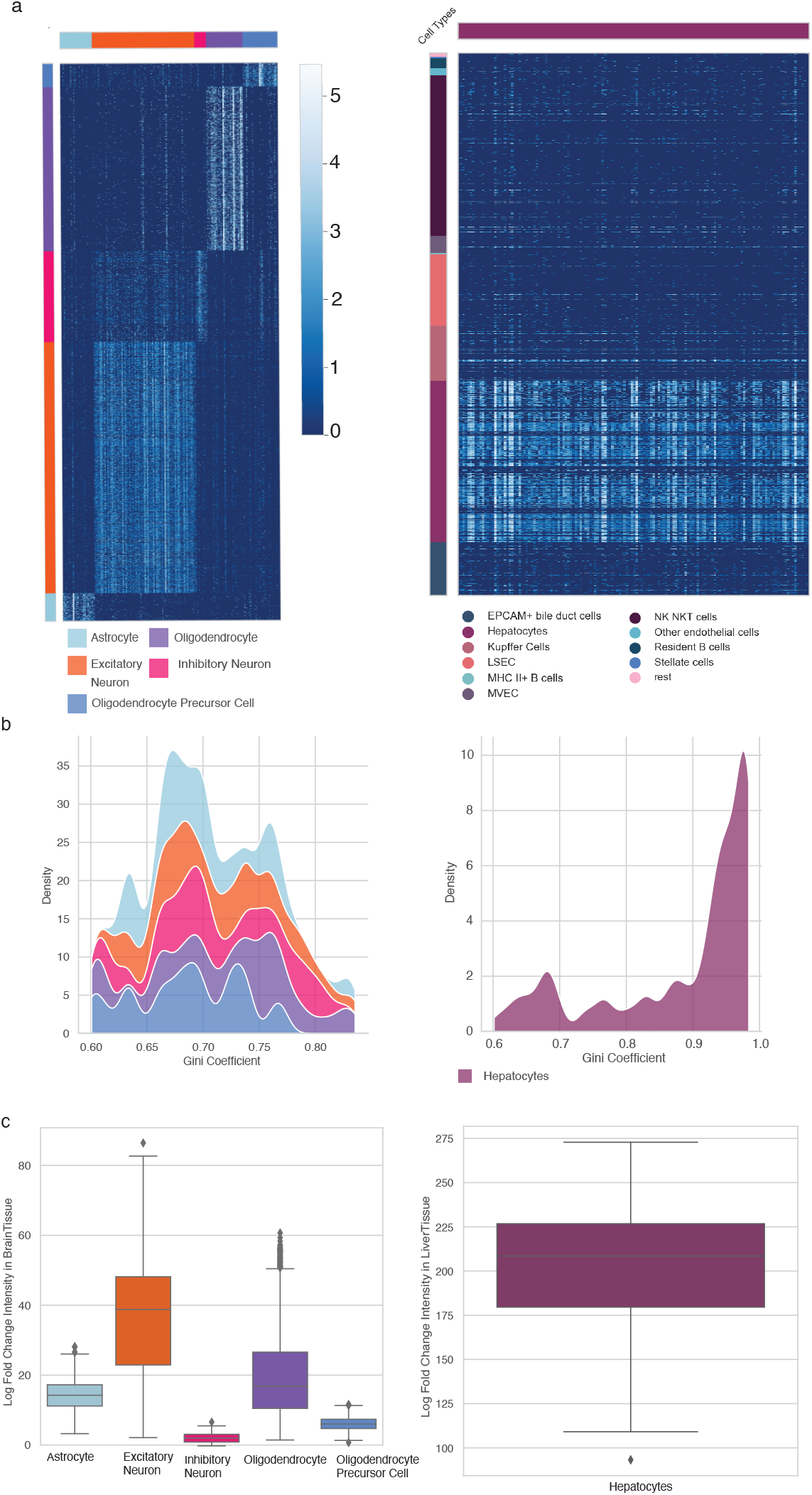
Establishing gene profile cell type specificity in context of the whole human body using single cell and bulk RNA-seq data. (a) Single cell heatmaps for gene cell type profiles within the corresponding tissue cell atlas, demonstrating that a cell type specific profile is unique to a given cell type across those within a given tissue. Columns denote marker genes for a given cell type; rows indicate individual cells. Heat mapped values are log-transformed counts-per-ten thousand. (b) Gini coefficient density plot for genes in cell type profiles derived from brain and liver single cell atlases using HPA NX counts. The area under the curve for a given cell type sums to one. (c) Log fold change in bulk RNA-seq data of a given cell type profile, demonstrating that the predominant expression of the cell type signature in its native tissue is highest relative to other non-native tissues. Values are the log-fold change of the signature score of a given cell type profile in the native tissue (indicated by the y-axis) to the mean expression in the remaining non-native tissues; this is the log ratio of the sum of log-transformed counts per ten-thousand balues. Box plot: horizontal line, median; lower hinge, 25th percentile; upper hinge, 75th percentile; whiskers span the 1.5 interquartile range; points outside the whiskers indicate outliers.

**Fig S9.**
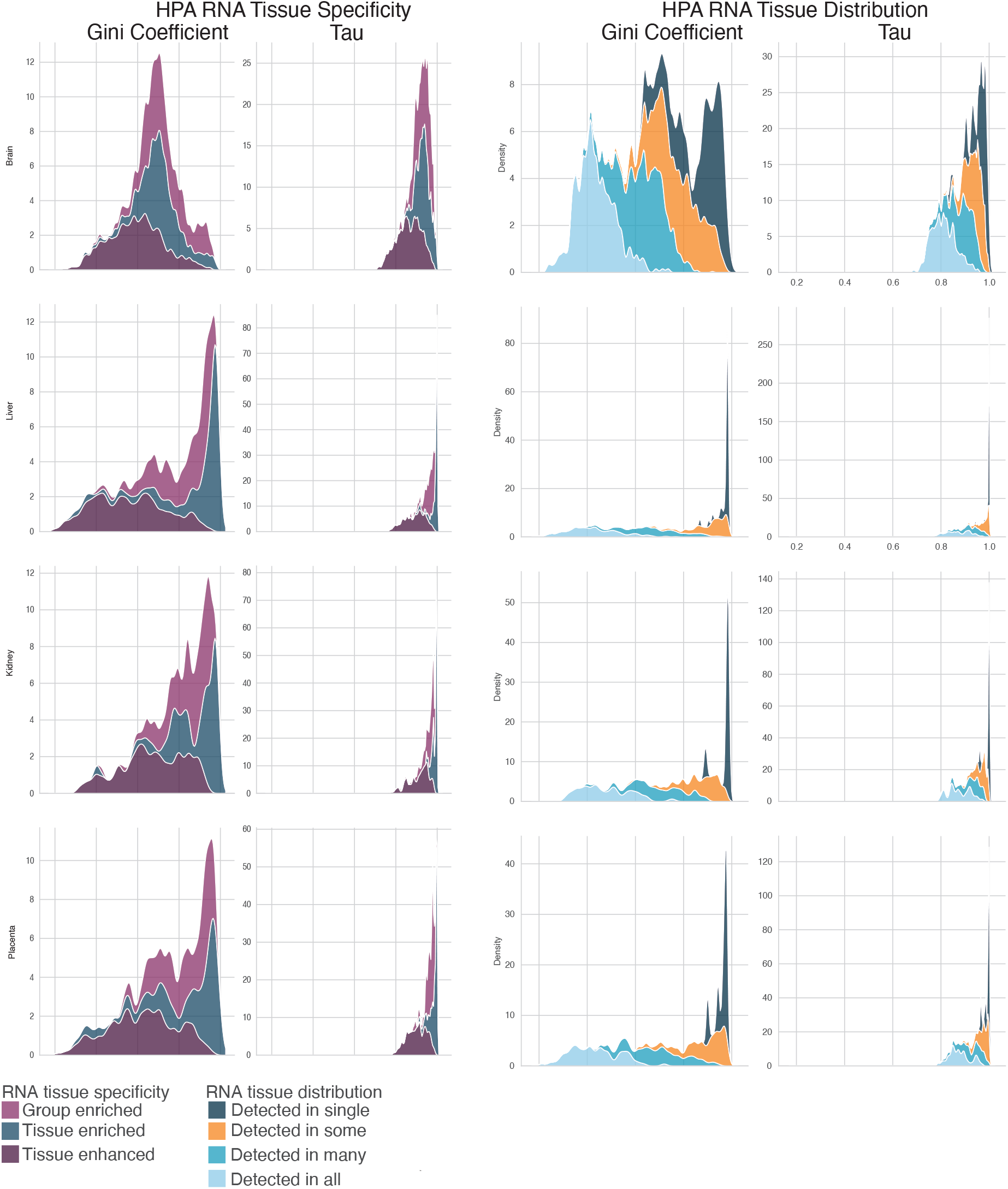
Distribution of Gini coefficient and Tau for all genes denoted by HPA as specific to the brain, liver, placenta, and kidney.

**Fig. S10.**
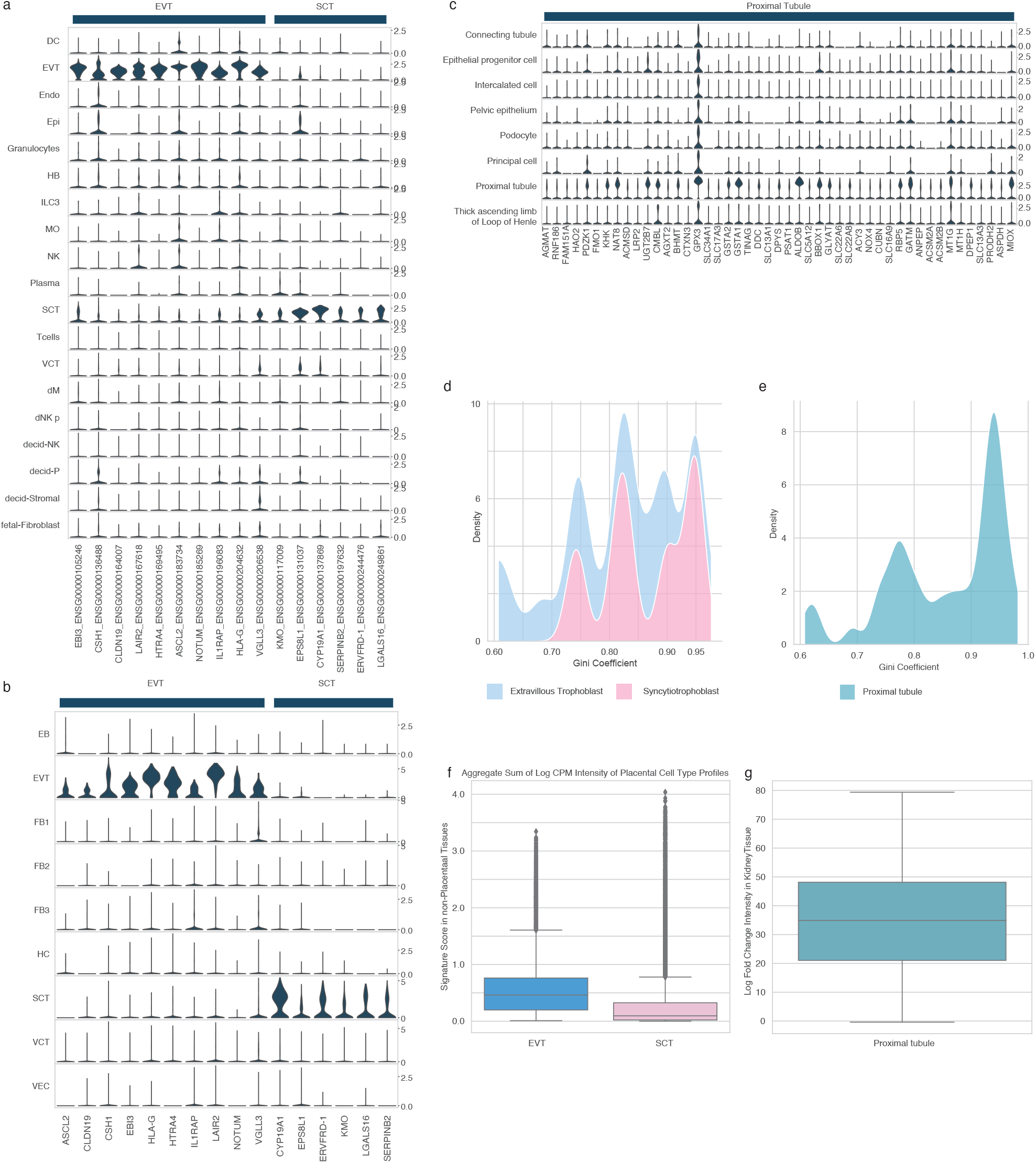
Comprehensive placental and renal cell type gene profile specificity at single cell and whole body resolution. For box plots in f, g : horizontal line, median; lower hinge, 25th percentile; upper hinge, 75th percentile; whiskers span the 1.5 interquartile range; points outside whiskers indicate outliers. (a) Violin plot of derived syncytiotrophoblast and extravillous trophoblast gene profiles from Vento-Tormo et al. (b) Violin plot of derived syncytiotrophoblast and extravillous trophoblast gene profiles from Suryawanshi et al. (c) Violin plot of derived proximal tubule gene profile (d) Gini coefficient distribution for placental trophoblast cell types in (a) and (b) (e) Gini coefficient distribution for renal cell type in (c) (f) Distribution of placental trophoblast signature scores across all GTEx tissues. Note: given that the placenta is not in GTEx, the box plots correspond to the distribution of signature scores across non-placental tissues (sum of log-transformed counts-per-ten thousand). (g) Log-fold change of renal cell type signature score in GTEx Kidney Cortex/Medulla samples relative to the mean non-kidney signature score, demonstrating that the predominant expression of the cell type signature in its native tissue is highest relative to other non-native tissues. Values are the log ratio of the sum of log-transformed counts per ten-thousand gene expression values.

**Fig. S11.**
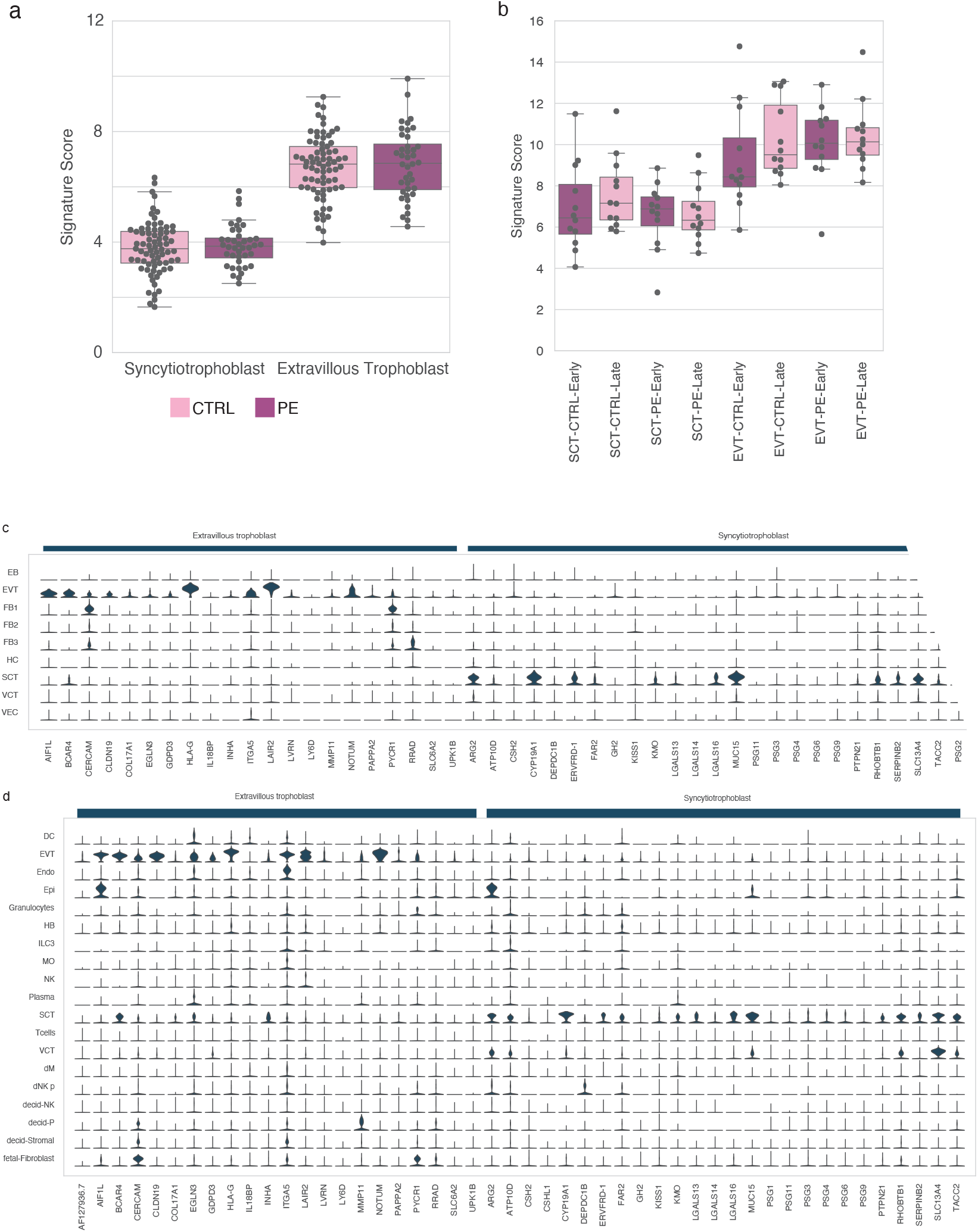
Expression distribution of Tsang et al trophoblast gene profiles and trophoblast signature scoring in preeclampsia cfRNA from (a) iPEC (mothers with no complications, n = 73; mothers with preeclampsia, n = 40) and (b) PEARL-PEC (n = 12 for each early/late-onset PE cohorts and gestationally-age matched healthy controls). Box plot: horizontal line, median; lower hinge, 25^th^ percentile; upper hinge, 75^th^ percentile; whiskers span the 1.5 interquartile range; points outside the whiskers indicate outliers. Stacked violin plot of the genes comprising the extravillous trophoblast and syncytiotrophoblast gene profiles from Tsang et al. intersecting with the measured genes in (c) Suryawanshi et al and (d) Vento Tormo et al, reflecting the expression distribution across placental cell types.

**Fig. S12.**
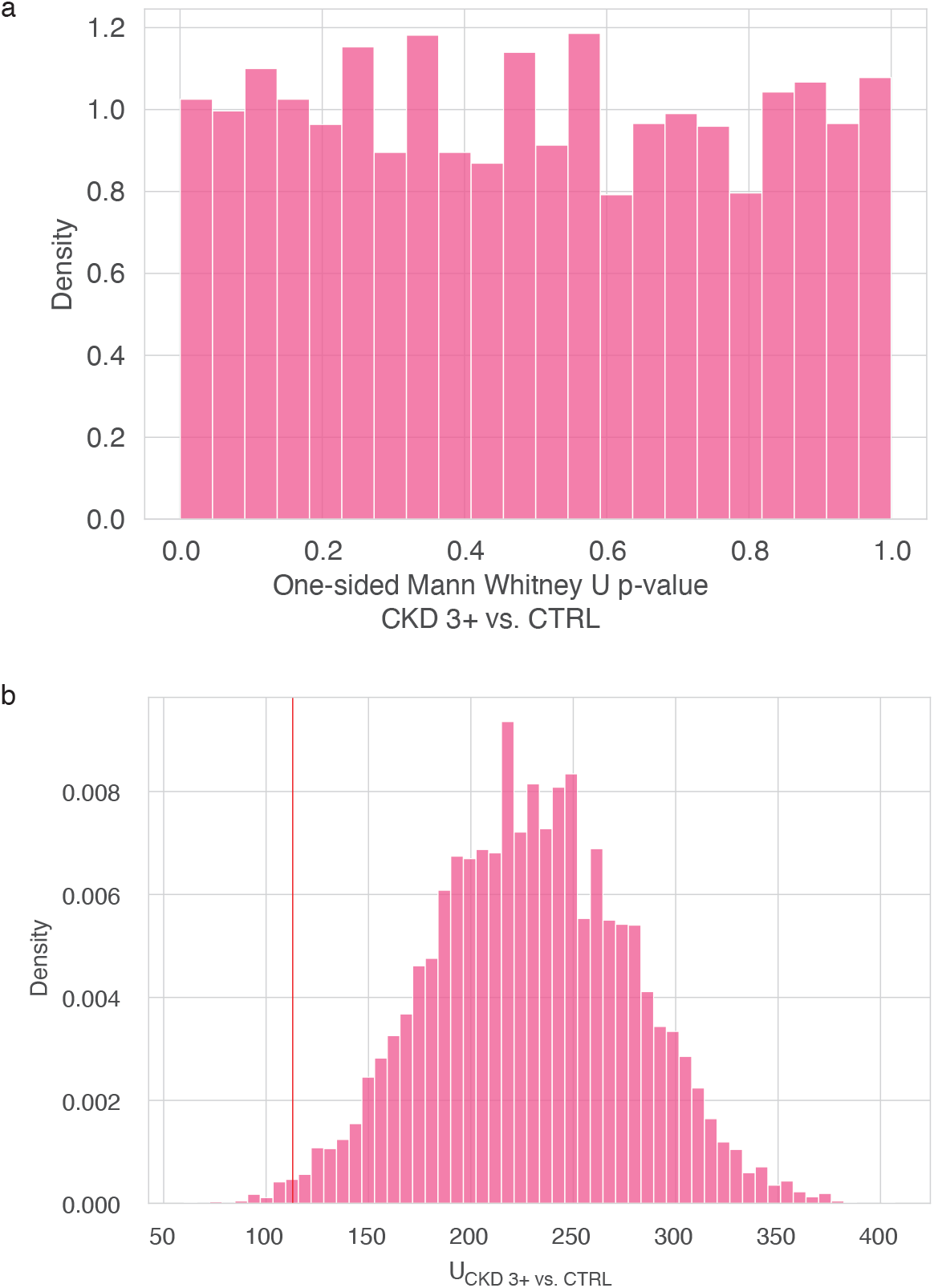
Assessment of p-value calibration for the proximal tubule signature score in CKD stage 3+ vs healthy samples following one-sided Mann Whitney U in 10,000 trial permutation test with an alternative hypothesis that the PT signature in healthy is greater than CKD (a) Distribution of p-values; distribution is uniform, as expected under the null. (b) distribution of U values; vertical line denotes the actual comparison; we observe a test-statistic as or more extreme than what was experimentally observed 0.63% of the time under the null. Taken together, this indicates the specificity of our signature score in discriminating between CKD and healthy patients.

**Fig S13.**
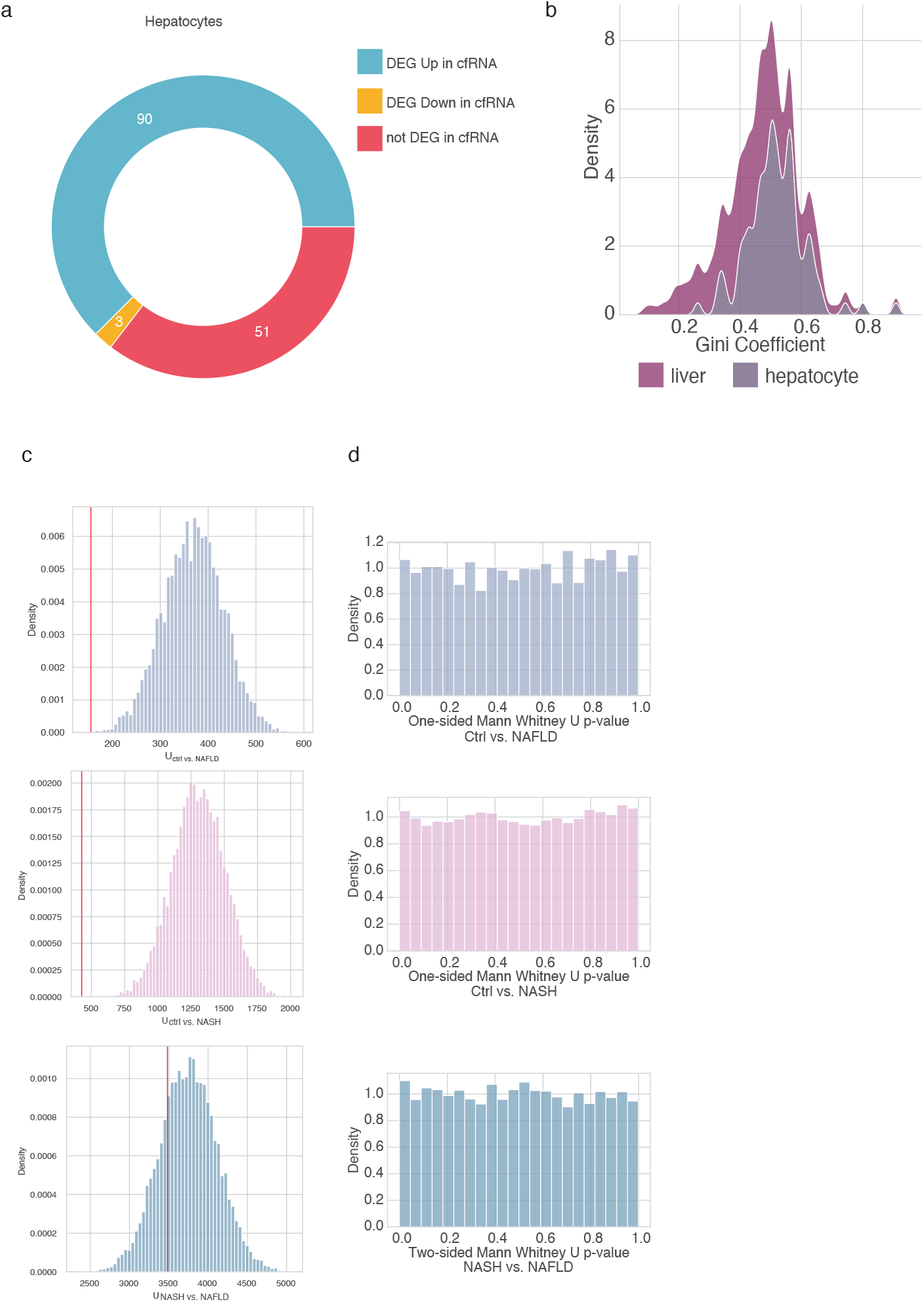
NAFLD/NASH DEG intersection with the hepatocyte gene profile and assessing their discriminatory power in signature scoring. (a) Donut plot reflecting the number of genes in the hepatocyte cell type gene profile that intersect with the reported NAFLD DEG^7^ (b) Density plot reflecting the Gini coefficient distribution corresponding to upregulated DEG in NAFLD^7^ that are liver or hepatocyte specific. The Gini coefficient is computed using the mean expression per liver cell type in Aizarani et al (Methods). Area under each curve sums to one. (c) Density of U values over 10,000 trial permutation test; red line indicates the U value corresponding to the experimental comparison reported in Fig. 2. (d) Density of p-values over 10,000 trial permutation test.

**Fig. S14.**
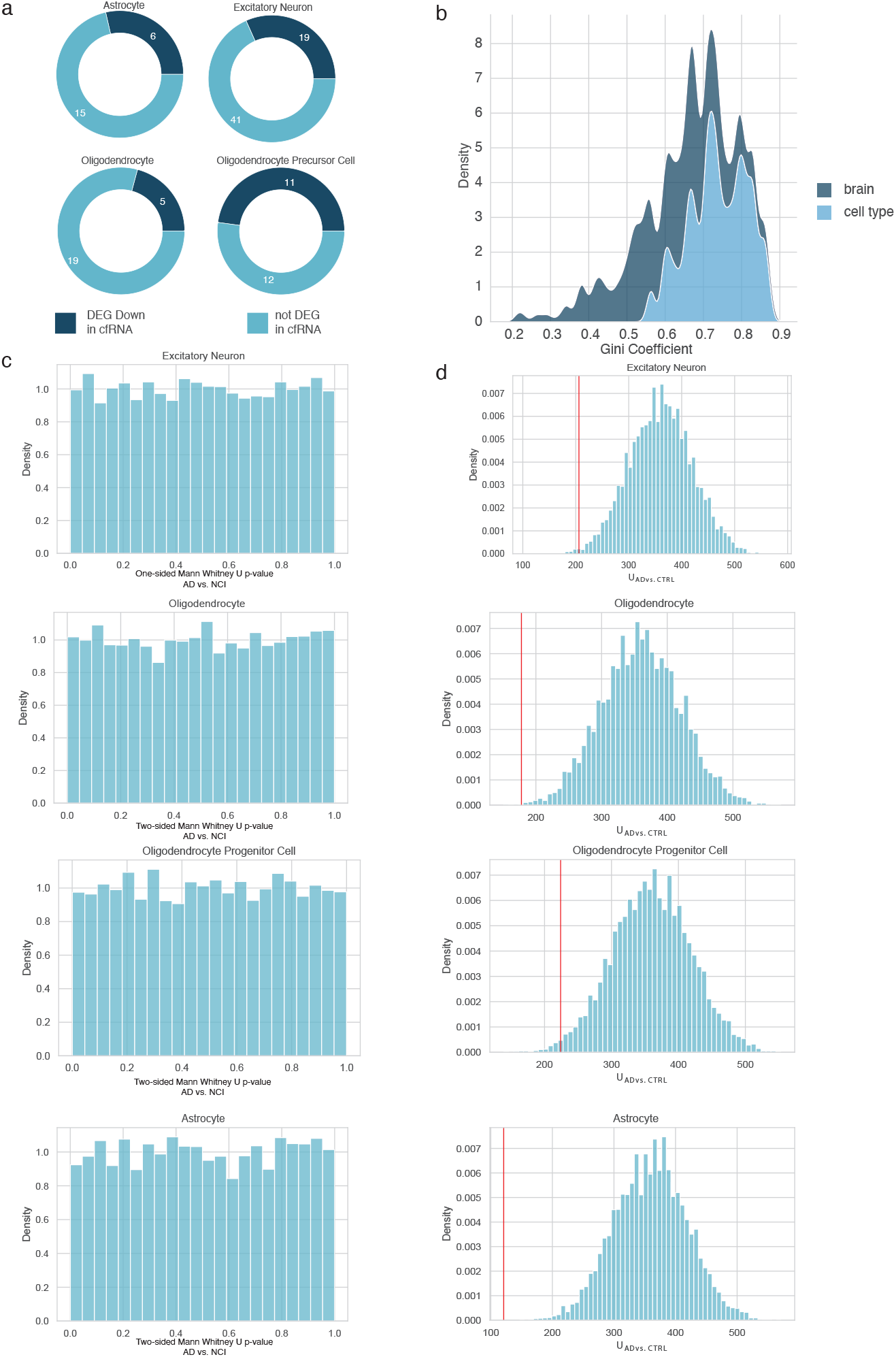
AD DEG intersection with neuronal/glial gene profiles and assessing their discriminatory power in signature scoring. (a) Donut plots reflecting the number of genes in the hepatocyte cell type gene profile that intersect with the reported NAFLD DEG^7^ (b) Density plot reflecting the Gini coefficient distribution corresponding to DEG in AD^7^ that are brain or brain cell type specific. The Gini coefficient is computed using the mean expression per brain cell type in the ‘Normal’ samples of Mathys et al (Methods). Area under each curve sums to one. (c) Density of p-values over 10,000 trial permutation test (d) Density of U values over 10,000 trial permutation test; red line indicates the U value corresponding to the experimental comparison reported in Fig. 2.

## References

1. Ibarra, A. et al. Non-invasive characterization of human bone marrow stimulation and reconstitution by cell-free messenger RNA sequencing. Nat. Commun. 11, 400 (2020).

2. Larson, M. H. et al. A comprehensive characterization of the cell-free transcriptome reveals tissue- and subtype-specific biomarkers for cancer detection. Nat. Commun. 12, 2357 (2021).

3. Ngo, T. T. M. et al. Noninvasive blood tests for fetal development predict gestational age and preterm delivery. Science 360, 1133–1136 (2018).

4. Koh, W. et al. Noninvasive in vivo monitoring of tissue-specific global gene expression in humans. Proc Natl Acad Sci USA 111, 7361–7366 (2014).

5. Munchel, S. et al. Circulating transcripts in maternal blood reflect a molecular signature of early-onset preeclampsia. Sci. Transl. Med. 12, (2020).

6. Toden, S. et al. Noninvasive characterization of Alzheimer’s disease by circulating, cell-free messenger RNA next-generation sequencing. Sci. Adv. 6, (2020).

7. Chalasani, N. et al. Noninvasive stratification of nonalcoholic fatty liver disease by whole transcriptome cell-free mRNA characterization. Am. J. Physiol. Gastrointest. Liver Physiol. 320, G439–G449 (2021).

8. Ngo, T. T. M., Moufarrej, M. N. & Rasmussen, M. L. H. Noninvasive blood tests for fetal development predict gestational age and preterm delivery. science.sciencemag.org.

9. Klatt, E. C. Robbins & Cotran Atlas of Pathology. (Elsevier, 2021).

10. Franzén, O., Gan, L.-M. & Björkegren, J. L. M. PanglaoDB: a web server for exploration of mouse and human single-cell RNA sequencing data. Database (Oxford) 2019, (2019).

11. Newman, A. M. et al. Robust enumeration of cell subsets from tissue expression profiles. Nat. Methods 12, 453–457 (2015).

12. Newman, A. M. et al. Determining cell type abundance and expression from bulk tissues with digital cytometry. Nat. Biotechnol. 37, 773–782 (2019).

13. The Tabula Sapiens Consortium & Quake, S. R. The Tabula Sapiens: a single cell transcriptomic atlas of multiple organs from individual human donors. BioRxiv (2021) doi:10.1101/2021.07.19.452956.

14. Sadeh, R. et al. ChIP-seq of plasma cell-free nucleosomes identifies gene expression programs of the cells of origin. Nat. Biotechnol. 39, 586–598 (2021).

15. Uhlen, M. et al. A genome-wide transcriptomic analysis of protein-coding genes in human blood cells. Science 366, (2019).

16. András, I. E. & Toborek, M. Extracellular vesicles of the blood-brain barrier. Tissue Barriers 4, e1131804 (2016).

17. Abbott, N. J. Inflammatory mediators and modulation of blood-brain barrier permeability. Cell. Mol. Neurobiol. 20, 131–147 (2000).

18. Ganong, W. F. Circumventricular organs: definition and role in the regulation of endocrine and autonomic function. Clin. Exp. Pharmacol. Physiol. 27, 422–427 (2000).

19. Suryawanshi, H. et al. A single-cell survey of the human first-trimester placenta and decidua. Sci. Adv. 4, eaau4788 (2018).

20. Vento-Tormo, R. et al. Single-cell reconstruction of the early maternal-fetal interface in humans. Nature 563, 347–353 (2018).

21. Stewart, B. J. et al. Spatiotemporal immune zonation of the human kidney. Science 365, 1461–1466 (2019).

22. Aizarani, N. et al. A human liver cell atlas reveals heterogeneity and epithelial progenitors. Nature 572, 199–204 (2019).

23. Kaufmann, P., Black, S. & Huppertz, B. Endovascular trophoblast invasion: implications for the pathogenesis of intrauterine growth retardation and preeclampsia. Biol. Reprod. 69, 1–7 (2003).

24. Tsang, J. C. H. et al. Integrative single-cell and cell-free plasma RNA transcriptomics elucidates placental cellular dynamics. Proc Natl Acad Sci USA 114, E7786–E7795 (2017).

25. Nakhoul, N. & Batuman, V. Role of proximal tubules in the pathogenesis of kidney disease. Contrib. Nephrol. 169, 37–50 (2011).

26. Chevalier, R. L. The proximal tubule is the primary target of injury and progression of kidney disease: role of the glomerulotubular junction. Am. J. Physiol. Renal Physiol. 311, F145–61 (2016).

27. Feldstein, A. E. & Gores, G. J. Apoptosis in alcoholic and nonalcoholic steatohepatitis. Front. Biosci. 10, 3093–3099 (2005).

28. Mathys, H. et al. Single-cell transcriptomic analysis of Alzheimer’s disease. Nature 570, 332–337 (2019).

29. Grubman, A. et al. A single-cell atlas of entorhinal cortex from individuals with Alzheimer’s disease reveals cell-type-specific gene expression regulation. Nat. Neurosci. 22, 2087–2097 (2019).

30. Dhillon, P. et al. The Nuclear Receptor ESRRA Protects from Kidney Disease by Coupling Metabolism and Differentiation. Cell Metab. 33, 379–394.e8 (2021).

31. Schelling, J. R. Tubular atrophy in the pathogenesis of chronic kidney disease progression. Pediatr. Nephrol. 31, 693–706 (2016).

32. Meex, R. C. R. & Watt, M. J. Hepatokines: linking nonalcoholic fatty liver disease and insulin resistance. Nat. Rev. Endocrinol. 13, 509–520 (2017).

33. McCall, M. A. et al. Targeted deletion in astrocyte intermediate filament (Gfap) alters neuronal physiology. Proc Natl Acad Sci USA 93, 6361–6366 (1996).

34. Lytton, J. Na+/Ca2+ exchangers: three mammalian gene families control Ca2+ transport. Biochem. J. 406, 365–382 (2007).

35. Friedman, L. G. et al. Cadherin-8 expression, synaptic localization, and molecular control of neuronal form in prefrontal corticostriatal circuits. J. Comp. Neurol. 523, 75–92 (2015).

36. Arlotta, P. et al. Neuronal subtype-specific genes that control corticospinal motor neuron development in vivo. Neuron 45, 207–221 (2005).

37. Shigemoto, R., Nakanishi, S. & Mizuno, N. Distribution of the mRNA for a metabotropic glutamate receptor (mGluR1) in the central nervous system: an in situ hybridization study in adult and developing rat. J. Comp. Neurol. 322, 121–135 (1992).

38. Zhou, Q., Choi, G. & Anderson, D. J. The bHLH transcription factor Olig2 promotes oligodendrocyte differentiation in collaboration with Nkx2.2. Neuron 31, 791–807 (2001).

39. Nielsen, J. A., Berndt, J. A., Hudson, L. D. & Armstrong, R. C. Myelin transcription factor 1 (Myt1) modulates the proliferation and differentiation of oligodendrocyte lineage cells. Mol. Cell. Neurosci. 25, 111–123 (2004).

40. Ichihara-Tanaka, K., Oohira, A., Rumsby, M. & Muramatsu, T. Neuroglycan C is a novel midkine receptor involved in process elongation of oligodendroglial precursor-like cells. J. Biol. Chem. 281, 30857–30864 (2006).

41. Levine, J. M., Reynolds, R. & Fawcett, J. W. The oligodendrocyte precursor cell in health and disease. Trends Neurosci. 24, 39–47 (2001).

42. Liddelow, S. A. et al. Neurotoxic reactive astrocytes are induced by activated microglia. Nature 541, 481–487 (2017).

43. Roadmap Epigenomics Consortium et al. Integrative analysis of 111 reference human epigenomes. Nature 518, 317–330 (2015).

44. Moufarrej, M. N., Wong, R. J., Shaw, G. M., Stevenson, D. K. & Quake, S. R. Investigating Pregnancy and Its Complications Using Circulating Cell-Free RNA in Women’s Blood During Gestation. Front. Pediatr. 8, 605219 (2020).

45. Pan, W. Development of diagnostic methods using cell-free nucleic acids. (Stanford University, 2016).

46. Robinson, M. D. & Oshlack, A. A scaling normalization method for differential expression analysis of RNA-seq data. Genome Biol. 11, R25 (2010).

47. Wolf, F. A., Angerer, P. & Theis, F. J. SCANPY: large-scale single-cell gene expression data analysis. Genome Biol. 19, 15 (2018).

48. Shen-Orr, S. S., Tibshirani, R. & Butte, A. J. Gene expression deconvolution in linear space. Nat. Methods 9, 9–9 (2011).

49. Chang, C.-C. & Lin, C.-J. Training nu-support vector regression: theory and algorithms. Neural Comput. 14, 1959–1977 (2002).

50. Zhong, Y. & Liu, Z. Gene expression deconvolution in linear space. Nat. Methods 9, 8–9; author reply 9 (2012).

51. Pedregosa, F. et al. Scikit-learn: Machine Learning in Python. Journal of Machine Learning Research 12, 2825–2830 (2011).

52. Qiao, W. et al. PERT: a method for expression deconvolution of human blood samples from varied microenvironmental and developmental conditions. PLoS Comput. Biol. 8, e1002838 (2012).

53. Kryuchkova-Mostacci, N. & Robinson-Rechavi, M. A benchmark of gene expression tissue-specificity metrics. Brief. Bioinformatics 18, 205–214 (2017).

54. van Rossum, D. & Hanisch, U.-K. Microglia. Metab. Brain Dis. 19, 393–411 (2004).

